# Transposons played a major role in the diversification between the closely related almond (*Prunus dulcis*) and peach (*P. persica*) genomes: Results from the almond genome sequence

**DOI:** 10.1101/662676

**Authors:** Tyler Alioto, Konstantinos Alexiou, Amélie Bardil, Fabio Barteri, Raúl Castanera, Fernando Cruz, Amit Dhingra, Henri Duval, Ángel Fernández i Martí, Leonor Frias, Beatriz Galán, José L. Garcia, Werner Howad, Jèssica Gómez Garrido, Marta Gut, Irene Julca, Jordi Morata, Pere Puigdomènech, Paolo Ribeca, María José Rubio Cabetas, Anna Vlasova, Michelle Wirthensohn, Jordi Garcia-Mas, Toni Gabaldón, Josep M. Casacuberta, Pere Arús

## Abstract

Combining both short and long-read sequencing, we have estimated the almond *Prunus dulcis* cv. Texas genome size in 235 Mbp and assembled 227.6 Mb of its sequence. The highly heterozygous compact genome of Texas comprises eight chromosomes, to which we have anchored over 91% of the assembly. We annotated 27,042 protein-coding genes and 6,800 non-coding transcripts. High levels of genetic variability were characterized after resequencing a collection of ten almond accessions. Phylogenomic comparison with the genomes of 16 other close and distant species allowed estimating that almond and peach diverged around 5.88 Mya. Comparison between peach and almond genomes confirmed the high synteny between these close relatives, but also revealed high numbers of presence-absence variants, many attributable to the movement of transposable elements (TEs). The number and distribution of TEs between peach and almond was similar, but the history of TE movement was distinct, with peach having a larger proportion of recent transpositions and almond preserving a higher level of polymorphism in the older TEs. When focusing on specific genes involved in key characters such as the bitter vs. sweet kernel taste and the formation of a fleshy mesocarp, we found that for one gene associated with the biosynthesis of amygdalin that confers the bitter kernel taste, several TEs were inserted in its vicinity only in sweet almond cultivars but not in bitter cultivars and *Prunus* bitter kernel relatives, including *P. webbii*, *P. mume*, and other species like peach and cherry. TE insertions likely to produce affects in the expression of six more genes involved in the formation of the fleshy mesocarp were also identified. Altogether, our results suggest a key role of TEs in the recent history and diversification of almond with respect to peach.

## Introduction

Almond, *Prunus dulcis* (Miller) D.A. Webb (syn. *P. amygdalus* Batsch), is a rosaceous tree species cultivated for its seeds, which has a diploid (2n=*2x*=*16*) and compact genome (∼250 Mbp). The *Prunus* genus comprises a group of approximately 200 species, some of which of high economic value, such as the stone fruit (peach, apricot, cherry and plum) and almond. The high level of genomic resemblance and synteny in the species of this genus (Dirlewanger et al. 2004) enables producing hybrids between some of them that are often fertile.

Humans used almond as food long before the advent of agriculture, and the oldest remains have been found in Israel, dating from 19,000 years ago (Kislev et al. 1992), although its domestication occurred probably 14,000 years later (Spiegel-Roy 1976). The origin of the almond tree is not well established, with its closest wild relatives living in Central and Western Asia, stretching from the Himalayas to the eastern Mediterranean basin (Yazbek and Al-Zein, 2014). Based on the distribution of the cultivated species, two alternative hypotheses place the domestication site of almond in the Levant (Browicz and Zohary, 1996) or in central Asia (Ladizinsky 1999). Diamond (1997) proposed almond as an example of simple domestication, where a dominant mutation at a single gene conferring sweet taste to the otherwise bitter and toxic kernel would result in an edible and cultivable crop. This gene, sweet kernel *Sk*/*sk*, was initially described by Heppner (1923) and later mapped to the central region of chromosome 5 (Sánchez-Pérez et al. 2007), although the causal DNA sequence remains to be identified. The closest relatives of almond are within the Amygdalus subgenus, encompassing peach [*P. persica* (L.) Batsch] and a group of 25 wild species (Yazbek and Al-Zein, 2014). Peach and almond hybrids are fertile. In fact, peach was proposed by Darwin (1868) as a possible direct derivative of almond with a fleshy, non-dehiscent and juicy mesocarp. However, molecular phylogenetics has identified a clear separation between peach and almond consistent with their geographic origin and distribution: peach and its closest relatives are native to China and eastern Asia, whereas almond and its wild relatives are native to central and western Asia (Delplancke et al. 2016).

The genome sequences of some *Prunus* species are available, including the high-quality genome of peach (Verde et al. 2013), and those of sweet cherry (*P. avium* L.) (Shirasawa et al. 2017), mume (*P. mume* L.), a relative of apricot (Zhang et al. 2012), and *P. yedoensis*, a wild cherry tree (Baek et al. 2018), the latter two used for ornamental purposes. In this paper, we present the whole genome sequence of almond cv. Texas, a self-incompatible and highly heterozygous genotype that was obtained in the United States from materials imported from Western Europe. Texas (also called Texas Prolific and Mission) was bred at Houston, Texas (US), as a seedling of French cultivar Languedoc (Wickson 1910), and became one of the leading cultivars of California in the last century along with ‘Nonpareil’ (Kester et al. 1991). Texas was also one of the parents – the other was peach cv. Earlygold – of the interspecific progeny used for the construction of the reference linkage map of *Prunus* (Joobeur et al. 1998). In this paper we describe the genome sequence of Texas almond and compare it with other sequenced genomes, including that of its close relative peach. We found that, in addition to other aspects of diversity between these genomes already reported (Velasco et al. 2016; Yu et al. 2018), transposable elements (TEs) played a key role in their short-term diversification.

## Methods

### DNA extraction, sequencing and K-mer analysis

Fresh young leaves of the Texas almond were ground in liquid nitrogen to a fine powder, and 100 mg of ground leaves were used for DNA extraction using the DNeasy® Plant Mini Kit (Qiagen, Valencia) according to the instructions of the manufacturer. The same method was employed for the extraction of ten additional almond cultivars (Aï, Belle d’Aurons, Cristomorto, Desmayo largueta, Falsa Barese, Genco, Marcona, Nonpareil, Ripon and Vivot), eleven peach cultivars (Armking, Belbinette, BigTop, Blanvio, Catherine, Earlygold, Flatmoon, Nectalady, Platurno, Sweetdream and Tiffany) and one accession of the wild almond *P. webbii* (R755). For sequencing with Oxford Nanopore Technologies (ONT) nanopore-based MinION sequencer, high molecular weight DNA from Texas almond was extracted with the method described by Mayjonade et al. (2016). For Texas, whole genome shotgun sequencing was performed using the Illumina HiSeq2500 and MiSeq sequencing instruments. The standard Illumina protocol with minor modifications was followed for the creation of short-insert paired-end libraries (Illumina Inc., Cat. # PE-930-1001). In brief, three libraries were generated from >2.0 μg of genomic DNA each. For one library the DNA was amplified by PCR while the other two libraries (and majority of sequence), the DNA was not amplified in order to reduce GC-bias. Then the DNAN was sheared on a Covaris™ E220, the fragmented DNA was size selected on an agarose gel to obtain three PE libraries with incremental insert sizes of 263 bp, 317 bp and 354 bp. The fragments were end-repaired, adenylated and ligated to Illumina indexed paired-end adaptors. The PE libraries were run on the of Illumina HiSeq2000 platform in 2×101 paired end mode according to standard Illumina operation procedures. Primary data analysis was carried out with the standard Illumina pipeline (HCS 2.0.12.0, RTA 1.17.21.3). A total of 97 Gb of raw sequence (>350x coverage) were produced. Post-processing of sequence reads involved detection and trimming of Illumina adapter sequence with cutadapt, quality trimming with trim_galore, and paired end overlap detection and merging with FLASH.

For the almond cultivars other than Texas and the peach and *P.webbii* accessions, we developed PE libraries of insert size 263 and Illumina sequenced as described in the previous paragraph.

Mate pair (MP) libraries of Texas DNA (3.1 and 5.2 kb fragment sizes) were constructed according to the Nextera Mate Pair Preparation protocol, which leaves a linker of known sequence at the junction. The resulting libraries were run on the HiSeq2000 platform in 2×101bp read length runs. Post-processing of sequence reads involved detection and trimming of the Nextera linker sequence with cutadapt, and quality trimming with trim_galore.

A Texas fosmid library was constructed in the pNGS vector (Lucigen). Sixty-eight pools of ∼500 clones per pool were made, and the purified DNA used to construct short-insert paired-end libraries. TruSeq™DNA Sample Preparation Kit v2 (Illumina Inc.) and the KAPA Library Preparation kit (Kapa Biosystems) were used. Briefly, 2.0 µg of genomic DNA were sheared on a Covaris™ E220, size selected and concentrated using AMPure XP beads (Agencourt, Beckman Coulter) in order to reach the fragment size of about 250bp. The fragmented DNA was end-repaired, adenylated and ligated to Illumina specific indexed paired-end adaptors. All libraries were quantified by Library Quantification Kit (Kapa Biosystems). The pools were sequenced using TruSeq SBS Kit v3-HS (Illumina Inc.), in paired-end mode, 2×150bp (45 libraries) and 2×250bp (23 libraries), in a fraction of a sequencing lane of HiSeq2500 flowcell (Illumina Inc.) according to standard Illumina operation procedures.

Jellyfish v2.2.6 (Marçais & Kingsford 2011) was run on the PE300 library (insert size 317bp, 2×100nt reads) with the Canonical kmer (-C) option and a mer-size of 21. GenomeScope (Vurture et al. 2017) was then used to analyze the resulting k-mer distribution. ONT reads were error-corrected using Canu (v1.5) (Koren et al. 2017). Corrected trimmed reads were used for hybrid assembly, scaffolding and assembly correction.

### Genome assembly

Given the relatively small size of the genome, we had the opportunity to experiment with different assembly strategies:

1. Whole genome shotgun (WGS) assembly based on paired-end (PE) and mate-pair (MP) 100nt read Illumina libraries.
2. Fosmid pool sequencing (150-250nt PE reads) and assembly combined with the WGS data.
3. Long read (ONT nanopore reads) and short read (Illumina PE) WGS hybrid assembly.

The short read WGS assembly with AbySS v1.3.6 (Simpson et al. 2009) resulted in a fragmented assembly (N50=4867 bp) with inflated genome size (512 Mbp). Heterozygosity and repeats were clearly going to be a problem. Fosmid-pool sequencing and assembly as previously described (Cruz et al. 2016; Abascal et al. 2016) increased contiguity (N50=142 kb) and reduced the assembly size (238 Mbp); however, the resulting assembly exhibited high levels of discordance with the available genetic maps, as well as with the peach assembly (v2.0.1). The final strategy, which included additional nanopore sequencing, resulted in the best balance of contiguity and concordance with the genetic map.

MaSuRCA v3.2.3 (Zimin et al. 2013) was run with default parameters (no linking mates; Celera assembly of super-reads). The input were the two PE Illumina libraries of insert sizes 317bp and 354bp for a total of 285x coverage (Table S1) and the self-corrected ONT reads for a total of ∼20x coverage.

Redundans v0.13 (Pryszcz & Gabaldón 2016) was run on the non-deduplicated output of MaSuRCA (the step 9-terminator genome.ctg.fasta file totaling 470Mb with N50=53kb) using the --longreads option for scaffolding, reducing the size of the assembly to 248Mb, 211Mb of which was scaffolded further with the ONT reads.

A first round of corrections to the assembly was carried out using consistency with nanopore data as the main criteria. Nanopore reads were mapped to the assembly with NGM-LRv0.2.6 (Sedlazeck et al. 2018), the assembly was broken at regions of zero coverage and then re-scaffolded with SSPACE-LongRead v1.1 (Boetzer and Pirovano 2014). A second round of corrections was made utilizing collinearity with the (Texas almond × Earlygold peach; TxE) genetic F2 and BC1 (to Earlygold) maps (Donoso et al. 2015) as the main criterion, with break points guided by synteny with the peach genome and coverage of nanopore reads. Peach transcripts (annotation Pp2.01a) were mapped to the almond assembly with GMAP v2014-12-23 (Wu and Watanabe 2005). Marker sequences were mapped with BWA mem keeping those mappings with mapping quality ≥20 and identity ≥90%. The broken assembly was again scaffolded with SSPACE-LR. The assembly at this stage had a contig N50 of 99kb and scaffold N50 of 151kb.

A third round of corrections was performed with improvements in mapping and break detection. First, peach transcripts were mapped only in the sense direction, and second, discrepant marker mappings were screened for mapping artifacts. Moreover, Sniffles v1.0.11 (Sedlazeck et al. 2018) was used for structural variant detection. Additional breaks were made and duplicate sequences were also detected. In the end, we were able to merge 30Mb with minimus2 from AMOS v3.1.0 (Sommer et al. 2007; Treangen et al. 2011) and remaining duplicate sequence (>99% identical, >5kb) was manually reviewed. Overlapping regions were joined into new longer contigs using nanopore read mappings to confirm new joins. Also at this stage, putative chloroplast sequence was identified by coverage and homology and set aside.

Finally, the assembly was anchored to pseudomolecules using both the TxE genetic map and synteny with the peach genome using ALLMAPS (jcvi-0.7.3) (Tang et al. 2015), with more weight given to the map marker order. Remaining conflicts were resolved manually. ALLMAPS uses a genetic algorithm for placing and orienting scaffolds, and sometimes it does not converge completely on the optimal solution, even with a large number of generations. Thus, we had to manually review and fix the order and orientation of some scaffolds which still exhibited discordance with either the genetic map or synteny with peach. Further improvement to the assembly was made by joining adjacent scaffolds if they could be linked together with split nanopore read mappings. A few additional overlaps were also detected in this fashion and longer contigs were constructed.

Assembly completeness was estimated in two ways. First, gene completeness was determined by running BUSCO v3.0.2 (Simão et al. 2015) using the embryophyta_odb9 database comprising 1,440 single-copy plant orthologous groups (BUSCOs). Second, a pairwise comparison of k-mers present in both input reads and the assembly was performed using KAT (Mapleson et al. 2017) using all WGS PE Illumina reads and a k-mer length of 27 (Figure S1).

### Comparison of the *P. dulcis* anchored assembly to the linkage map and peach v2.0 a1 genome sequence

The almond assembly was compared to the TxE linkage map that contains 1,833 SNP markers (Donoso et al. 2015). Markers were mapped onto the almond pseudomolecule-based assembly using BLAST and coordinate data of both almond and peach were used as input in MapChart software (Voorrips *et* al., 2002) for representing graphically the comparison between the two species. Genetic and physical distance of SNP markers from the TxE population were used for calculating the recombination rate across the pseudomolecules of the almond assembly. In-house python scripts were used for generating the corresponding graphs.

The peach genome sequence and annotation data were downloaded from GDR ftp site (ftp://ftp.bioinfo.wsu.edu/species/Prunus_persica/Prunus_persica-genome.v2.0.a1/). Synteny between almond with peach genomes was assessed using the SyMap software v4.2 (Soderlund et al., 2006) with default parameters, except that the “min dot” parameter was set to 25.

### Annotation

The annotation of the *P. dulcis* genome assembly was performed by combining transcript alignments, protein alignments and ab initio gene predictions. A flowchart of the annotation process is shown in Figure S2

First, almond RNAseq reads were downloaded from NCBI with the accession number SRR1251980 and aligned to the genome with STAR (v-2.5.3a) (Dobin et al. 2013). Transcript models were subsequently generated using Stringtie (v1.0.4) (Pertea et al. 2015) and, along with the *P. persica* transcriptome (annotation Pp2.0a) and 4,509 almond ESTs downloaded from NCBI on July 2015, were assembled into a non-redundant set by PASA (v2.3.3) (Haas et al. 2008). The TransDecoder program, which is part of the PASA package, was run on the PASA assemblies to detect coding regions in the transcripts. Second, the complete Rosaceae proteome was downloaded from Uniprot on July 2015 and aligned to the genome using Exonerate (v2.4.7) (Slater and Birney, 2005). Third, *ab initio* gene predictions were performed on the repeat masked pdulcis26 assembly with three different programs: GeneID v1.4 (Alioto et al. 2018), Augustus v3.2.3 (Stanke et al. 2006) and GeneMark-ES v2.3e (Lomsadze et al. 2014) with and without incorporating evidence from the RNAseq data. Finally, all the data was combined into consensus CDS models using EvidenceModeler-1.1.1 (EVM) (Haas et al. 2008). Additionally, UTRs and alternative splicing forms were annotated through two rounds of PASA annotation updates.

The annotation of non-coding RNAs was produced by running the following steps. First, the program cmsearch v1.1 (Cui et al. 2016) from the INFERNAL package (Nawrocki and Eddy 2013) was run against the RFAM (Nawrocki et al. 2015) database of RNA families (v12.0). Also, tRNAscan-SE v1.23 (Lowe & Eddy 1997) was run to detect the transfer RNA genes present in the genome assembly. To annotate long non-coding RNAs (lncRNAs) we first selected PASA-assemblies that had not been included in the annotation of protein-coding genes. Those longer than 200bp and whose length was not covered at least in an 80% by a small ncRNA were incorporated into the ncRNA annotation as lncRNAs. The resulting transcripts were clustered into genes using shared splice sites or significant sequence overlap as criteria for designation as the same gene.

#### Functional annotation

Functional annotation was performed using an in-house pipeline that integrates several data sources to infer protein function based on sequence similarity to annotated sequences or/and presence of particular domains and sequence motifs. We used InterPro (Hunter et al. 2011), KEGG (Kanehisha et al. 2012), signalP (Petersen et al. 2011), and NCBI CDsearch (Manchler-Bauer et al. 2011) databases. InterProScan v.5.19-58 (Zdobnov et al. 2001) was used to scan though all available InterPro databases, including PANTHER, Pfam, TIGRFAM, HAMAP and SUPERFAMILY. Initial sequence similarity search was determined using BLASTP v.2.6.0+ against NCBI non-redundant (NR) collection of protein sequences (release 2018-08). KEGG orthology (KO) groups were assigned by KEGG Automatic Annotation Server (KAAS) (Moriya et al. 2007) using the bi-directional best hit (BBH) method against a representative gene set from 27 different species, which includes a core set of species for gene annotation and additional plant species from the Rosaceae family. KO identifiers were then used to retrieve the KEGG relevant functional annotation using the KEGG REST-based API service, KEGG release v.87.1.

To predict plant disease resistance genes, each protein was searched against a manually curated list of ‘reference’ R-genes with the DRAGO pipeline (Sanseverino et al. 2013). For each hit, classes were assigned based on combination of specific domains, such as TIR (Toll-Interleukin like Region), NBS (Nucleotide Binding site), LRR (Leucine-Rich Region), or coiled-coil domain. Putative transcription factor genes were predicted by using the Plant Transcriptional Factor database (Jin et al. 2017) v.4.0

#### Annotation and analysis of transposable elements in genome assemblies

The Ilumina paired-end reads corresponding to the resequencing of the almond and peach varieties described in section 2.1 have been trimmed with SKEWER (version 0.2.2, --mean-quality 25, --min 35) and aligned to their respective reference genome with BWA-aln/sampe^1^ (version 0.7.5, parameters: -t 6, -n 5, -o 1, -e 3) (Li and Durbin, 2009) and SAMTOOLS (version 0.1.18) (Li 2011). Bam files were later submitted to the package PINDEL (version 0.2.5, parameters: -T 4, -x 5, -r false, -t false, -A 35) (Ye et al. 2009) to identify deletions in samples. Transposable elements (TE) were annotated in *P. dulcis* and *P. persica* assemblies using TEdenovo and TEannot pipelines of the REPET package (Quesneville et al., 2005; Flutré et al, 2011) installed in the PiRATE virtual machine (Berthelier et al., 2018). Classification of TEdenovo consensus sequences at the order level was done with PASTEC (Hoede et al., 2014).

TE annotation for masking purposes was obtained using RepeatMasker (www.repeatmasker.com) with a reduced TE representative library. The TE representatives obtained using the TEdenovo pipeline library were screened for coding domains with hmmscan (HMMER 3.1b1, hmmer.org) against the PFAM database (Finn et al., 2016). TE representatives containing regions potentially coding for known domains of non-TE proteins usually found in multigene families (kinases, NB-ARC, LRR, TIR) or with an N content higher than 30% or a with length shorter than 200 nucleotides were discarded. Moreover, all TE representatives not categorized in one of the classical TE superfamilies (defined as “noCat” by TEdenovo) were also removed. In total, 661 representatives were removed from the library. The final library contains 6,898 TE representatives.

MITE-hunter (Han and Wessler, 2010) was run to detect potential MITE families. In order to complete the annotation the potential almond MITE families were combined with the *P. persica* family annotation available in PMITE database (families carrying TSD) (Chen et al. 2014). These sequences were grouped in clusters of 90% identity with cd-hit (Fu et al., 2012) to remove redundancy and produce a final library of family representatives. RepeatMasker (http://www.repeatmasker.org/) was run to annotate all regions having significant similarity to MITE families, and the results were filtered to retain only full-length elements (consensus length ± 20%). The same pipeline was used to identify MITEs in the *P. persica* assembly.

Long Terminal Repeat retrotransposon (LTR-retrotransposon) candidates were predicted by running LTRharvest (Ellinghaus et al. 2008) with default parameters. The internal conserved domains of these elements were identified using HMMER hmmscan (Johnson et al. 2010) and only coding elements were retained for further analyses. Elements displaying either a single hit on the genome, more than 10% of gaps or more than 50% of tandem repeats were filtered out. Classification of the remaining elements (hereafter referred as “coding LTR-retrotransposons”) into Copia and Gypsy superfamilies was performed based on the order of the internal coding domains, as defined by Xiong & Eickbush (1990). Elements lacking one or more domains were tagged as “incomplete”.

The LTR regions of every coding element were extracted and aligned with MUSCLE (Edgar et al. 2004). Kimura 2P distance of every aligned LTR pair was calculated and used to estimate insertion ages following the approach described in San Miguel et al. (1998), using a substitution rate of 10^−8^ nucleotides per site per year and a generation time of 10 years (Velasco et al. 2016).

The flanking sequences (500bp) of every coding LTR retrotransposon were extracted from *P. dulcis* and used as query for a BLASTn (Altschul et al. 1990) search (cut off E value < e-10) against the *P. persica* assembly, and vice-versa. Concordant mapped flanks were defined when both flanks of an element mapped in the same scaffold at a distance smaller than 25 kb. Every internal region between two concordant mapped flanks was aligned to the putative orthologous element using EMBOSS Needle (Rice et al. 2000). The two elements were considered orthologous if they could be aligned over 80% of their length with at least 80% of identity. To assess the orthology of MITE insertions the approach followed was as the one described for LTR retrotransposons except that the sequences flanking the insertions were mapped to the corresponding genome using BBmap (https://sourceforge.net/projects/bbmap/) instead of Blast.

In order to search for polymorphic LTR-retrotransposon and MITE insertions within or close to genes we used BEDTools (version 2.27.0) (Quinlan and Hall, 2010). Only TEs located within genes or at less than 1000 nt upstream of a gene were kept.

### Analysis of DNA methylation in almond and peach

After bisulfite conversion, DNA from *P. dulcis* (cv. Texas, almond) and *P. persica* (var. “Earlygold”, peach) young leaves was sequenced using Illumina (Hi-Seq, 2 × 101-nt read length). Raw reads were trimmed with TrimGalore! version 0.4.5 (http://www.bioinformatics.babraham.ac.uk/projects/trim_galore/). Low quality bases (Phred score < 20) were trimmed before adapter removal and reads with a length less than 20 were discarded. The total of trimmed reads was 82,073,678 and 94,040,426 in almond and peach, respectively. Trimmed reads of each species were mapped to their respective reference genome and methylation was analyzed using Bismark version 0.19.1 (Krueger and Andrews, 2011). The gene/TE methylation was analyzed with SeqMonk version 1.41 (http://www.bioinformatics.babraham.ac.uk/projects/seqmonk/). Only cytosine positions that had been sequenced at least three times were included.

### Analysis of the *CYP71AN24* locus in *Prunus* related species and almond varieties

A two Mb region of the *P. dulcis* genome containing the *CYP71AN24* gene was compared to the corresponding genome regions of *P. avium, P. mume, P. persica* using Mauve (Darling et al. 2004). Resequencing data from the *P. dulcis* varieties Texas, Marcona, Cristomorto, D05-187 (SRX245830) and S3067 (SRX245832), as well as from *P. webbii* (R755), were mapped to the almond reference genome using BWA aln/sampe (Li and Durbin, 2009)

### *P. dulcis* phylome reconstruction

The *P. dulcis* and *P. persica* phylomes – i.e. the complete collection of evolutionary histories of all encoded genes – were reconstructed using the PhylomeDB pipeline (Huerta-Cepas et al., 2011). In brief, for each protein-coding gene in the almond and peach genome we searched for homologs (Smith-Waterman Blast search, e-value cutoff < 1e-05, minimum contiguous overlap over the query sequence cutoff ≥50%) in a database containing the proteomes of 17 species with sequenced genomes representing most of the important plant families (Table S2). The most similar 150 homologues were aligned using three different programs MUSCLE (Edgar, 2004), MAFFT (Katoh et al., 2005) and KALIGN (Lassmann & Sonnhammer, 2005) in forward and reverse orientation. These six alignments were combined using M-COFFEE (Wallace et al., 2006), and trimmed with trimAl v.1.3 (Capella-Gutiérrez et al., 2009), using a consistency cut-off of 0.16667 and a gap threshold of 0.1. Phylogenetic trees were built using a Maximum Likelihood approach as implemented in PhyML v3.0 (Guindon & Gascuel, 2003) using the best fitting model among seven different ones (JTT, LG, WAG, Blosum62, MtREV, VT and Dayhoff). The model best fitting the data was determined comparing the likelihoods estimated on an initial Neighbor Joining tree topology and using the AIC criterion. In all cases we used four rate categories and inferred the fraction of invariant positions and rate parameters from the data. Then, these phylomes were filtered to remove the gene trees that contain proteins associated to transposon-related functional terms. All alignments and trees are available for browsing or download at PhylomeDB with the PhylomeID 406 (almond phylome) and 407 (peach phylome) (Huerta-Cepas et al., 2014; www.phylomedb.org).

Orthology and paralogy relationships were predicted based on phylogenetic evidence from the almond and peach phylomes. We used ETE v3 (Huerta-Cepas et al., 2010) to infer duplication and speciation relationships using a species overlap approach and a species overlap score of 0. The relative age of detected duplications was estimated using a phylostratigraphic approach that uses the information on which species diverged prior and after the duplication node (Huerta-Cepas & Gabaldón, 2011). Duplication frequencies at each node in the species tree were calculated by dividing the number of duplications mapped to a given node in the species tree by all the gene trees that contain that node. For this analysis, we excluded gene trees that contained large (more than 5 paralogs) species-specific expansions (expansions that contained more than five members). All orthology and paralogy relationships are available through PhylomeDB (Huerta-Cepas et al., 2014).

Gene Ontology (GO) term enrichment analysis was performed using FatiGO (Al-Shahrour et al., 2007). We compared three lists of proteins against all the other proteins encoded in the genome. The three lists were composed of the proteins involved in a duplication at the ancestral node of all *Prunus* species, the proteins specifically lost in almond, and the proteins specifically lost in peach.

The trimmed alignments of 262 genes that had single-copy orthologs in the 17 species considered were selected and concatenated. The final alignment containing 141,911 amino acid positions was used to reconstruct the maximum likelihood species tree with PhyML v3.1 (Guindon et al., 2010) using the LG amino acid substitution model, and 100 bootstrap replicates. Additionally, a super-tree was reconstructed using all trees in the phylome and a gene tree parsimony approach as implemented in duptree (Wehe et al. 2008).

Divergence dates were estimated on the topology derived from the Maximum Likelihood approach by using the Bayesian relaxed molecular clock approach as implemented in PhyloBayes v4.1c (Lartillot et al. 2013). An uncorrelated relaxed clock model was applied, and four fossil constraints specified to the most recent common ancestor: *Prunus* (47.8 Mya, Li et al. 2011), Rosaceae (98.25 Mya, Crepet and Nixon, 1996, Zhang et al. 2017), split between Fagales and Cucurbitales (84 Mya, Herendeen et al. 1995, Sims et al. 1998, Wikström et al. 2001), Eudicots (124 Mya, Hughes and McDougall, 1990).

These calibration constraints were used with soft bounds (Yang and Rannala, 2006) under a birth-death prior, and a prior on the root of the tree (183 Mya) (Bell et al., 2010). Two independent MCMC chains were run for 20,000 cycles, sampling posterior rates and dates every 10 cycles. The initial 25% were discarded as burn-in. Posterior estimates of divergence dates and associated 95% credibility intervals were then computed from the remaining samples of each chain.

### Resequencing of almond cultivars and almond-peach structural genome variability comparison

Genetic variability analysis was performed on ten almond traditional cultivars and one peach cultivar, Earlygold, that was used as outgroup. These accessions were resequenced using paired-end Illumina sequencing as described in section 2.1. Selection of the almond lines was based on their origin as representing a range of the major areas of production in Europe (France, Spain, Italy) and the USA, and morphological characteristics (shell hardness, bloom time and self-incompatibility) (Table S3). Earlygold is the peach cultivar that was used with Texas almond to create the linkage map that is used as the reference for *Prunus* (Dirlewanger et al. 2004).

Paired-end Illumina sequencing data from the almond varieties were trimmed (length ≥ 35bp, mean sliding window of 4bp phred quality score ≥ 20) using Trimmomatic (Bolger AM *et al*., 2014) and the output was quality checked using FastQC (https://www.bioinformatics.babraham.ac.uk/projects/fastqc/). Trimmed data were aligned against the almond assembly using the BWA-MEM algorithm v0.7.16a-r1181 (http://bio-bwa.sourceforge.net/bwa.shtml) with default parameters. After removal of unmapped reads and marking of PCR duplicates, variant calling was performed using Samtools v1.5 (Li 2011) with default parameters, except from the following: -q 10 -Q 20. Variant calling format (VCF) files were filtered by applying the following criteria: global quality ≥ 30, genotype quality ≥ 30, 8≤depth≤ 300, biallelic sites, 0.1≤MAF. Graphical representation of variant distributions was done with Circos (Krzywinski et al. 2009) in non-overlapping windows.

Large deletions between almond and peach resequencing data were identified by comparing with the almond assembly using Pindel (Ye et al. 2009) using default parameters and an insert size of 300bp for all the samples. Peach- and almond-specific deletions were obtained by selecting positions with at least 20 reads/cultivar supporting the event. We also removed deletions that overlapped with N-regions (±1000bp) in the almond genome. We considered that a position overlaps with a TE if at least one of the two elements, the deletion or the TE, had at least 80% of its sequence overlapping with the other element.

*P. dulcis* contigs were aligned to the *P. persica* reference genome using Nucmer from the Mummer3 package (Delcher et al. 2003). Assemblytics (Nattestad and Schatz, 2016) was used to filter the alignment and detect genome-wide variants with the following cut-offs: Unique sequence length required for considering an alignment = 10, 000 bp, minimum variant size= 20bp, maximum variant size= 25,000 bp. Structural variants were intersected with TE annotations of *P. dulcis* (insertions and repeat expansions) and *P. persica* (deletions and repeat contractions). A variant was considered to be TE-associated when at least 50% of its sequence was spanned by a TE.

## Results

### Sequence analysis and genome size estimation

A total of 97 Gb of paired-end Illumina sequence (>350x coverage) and 31.6 Gb of Illumina mate-pair sequence were produced. The yield per fosmid pool was on average 5.2 Gb (3.4 – 9.0 Gb) of raw sequence, corresponding to a sequencing depth of 250x (170-450x). Oxford Nanopore (ONT) sequencing produced 10.2 Gb of sequence with Q≥7.0 with a read N50 length of 7.3 kb. More than 82% of reads had an average Q score ≥ 10. Error correction with Canu yielded 4.5 Gb of corrected trimmed reads with a read N50 of 11.5 kb, which were used for assembly. A summary of sequencing library statistics is given in Table S1. By analyzing k-mer frequency (Jellyfish/Genomescope), the genome size was estimated to be 238 Mb. A plot of k-mer coverage is shown in Figure S3.

### Sequence assembly and annotation

We performed a hybrid assembly strategy combining Illumina and Oxford Nanopore sequence data. We collapsed the assembly into a haploid representation and performed a series of mis-assembly detection and correction steps before anchoring the assembly to eight pseudomolecules, the number of the haploid almond chromosome complement. The final assembly, *P. dulcis* Texas v2.0 (a.k.a. pdulcis26) totals 227.6 Mb (91.5% of which is anchored to the eight pseudomolecules) and has a contig and scaffold N50s of 103.9kb and 381.5kb, respectively (Table 1). This assembly is available at the Genome Database for the Rosaceae (GDR; https://www.rosaceae.org)

**Table 1.** Texas genome assembly and annotation statistics

|  |  |
| --- | --- |
| Assembly length | 227.6 Mb |
| Contig N50 | 103.9 kb |
| Scaffold N50 | 381.5 kb |
| Pseudomolecule N50 | 24.8 Mb |
| Percent anchored to pseudomolecules | 91.47% |
| BUSCO complete genes | 95.4% |
| BUSCO fragmented genes | 1.0% |
| BUSCO missing genes | 3.6% |
| Genomic GC content | 37.65% |
| Number of protein-coding genes | 27,042 |
| Median gene length (bp) | 2,315 |
| Number of transcripts | 33,119 |
| Number of exons | 144,681 |
| Number of coding exons | 136,958 |
| Coding GC content | 44.91% |
| Median intron length (bp) | 172 |
| Exons/transcript | 4.37 |
| Transcripts/gene | 1.22 |
| Multi-exonic transcripts | 81% |
| Gene density (kb/gene) | 8.41.42 |

The completeness of the assembly as determined by BUSCO analysis is 96.4%: 95.4% complete (89.4% unique and 6.0% duplicated), 1.0% fragmented and 3.6% missing BUSCOs. A k-mer analysis (Figure S3) comparing the coverage of k-mers extracted from the Illumina reads and their multiplicity in the final assembly confirms the BUSCO results, with a small fraction of missing unique sequence as well as some level of duplicated sequence, which can be attributed to the difficulties associated with assembling highly heterozygous genomes such as this one.

In total, we have annotated 27,042 protein-coding genes that produce 33,119 transcripts (1.22 transcripts per gene) and encode for 31,654 unique protein products. We were able to assign some type of functional annotation to 92% of them. The annotated transcripts contain 4.37 exons on average, with 81% of them being multi-exonic (Table 1). In addition, we annotated 6,800 non-coding transcripts, of which 3,604 and 3,196 are long and short non-coding RNA genes, respectively.

### Comparison of the anchored almond assembly with the linkage map and the peach sequence

Out of the 1,833 SNPs that comprise the TxE linkage map (Donoso et al. 2015), 1,609 (87.8%) mapped onto the almond assembly with single high quality hits (percentage identity ≥90% and marker coverage ≥ 90%) (Table S4), with 1,597 (93.4%) mapping onto the anchored assembly. Of the anchored SNPs, 1,578 aligned with the pseudomolecules and were syntenic and collinear to the eight chromosomes of peach. Only 19 SNPs had a different order on the assembly, 12 of which mapped onto a pseudomolecule or scaffold that was different from the linkage group of the TxE map (Table S4). Graphs generated with MapChart show the high synteny between the almond pseudomolecules and their corresponding linkage groups in the TxE genetic map. Minor discrepancies were observed for all the pseudomolecules except Pd05 (Figure S4; Table S4), which we attributed to contig reordering or minor misassemblies. Similarly, we observed high synteny and collinearity between the genome sequence of peach v2.0 a1 (Verde et al. 2017) and that of the almond Texas generated here (Figure S5). The comparison of physical vs. genetic distances of the eight pseudomolecules is presented in Figure S6. Regions of low recombination rates (horizontal fragments of the curve or fragments with low slope) usually coincide with pericentromeric regions and occurred at similar regions for every almond chromosome as in the peach genome (Verde et al. 2013).

### Transposable element landscape

Using the REPET pipeline we annotated the 38.21% of the almond genome as TE-related sequences (Table S5 and S6). The distribution of TEs along almond pseudochromosomes shows an inverse correlation with respect to the gene density, with TE-rich regions showing low gene density per chromosome, coinciding with pericentromeric regions, and lower TE densities in the gene-rich chromosomal arms (Figure 1A). The almond TE landscape was compared with that of peach. For that purpose, we annotated peach TEs with the same strategy and found very similar results: 37.60% of TE content (Table S5 and S6) and a comparable TE and gene distribution to that of almond chromosomes (Figure 1B).

**Figure 1.**
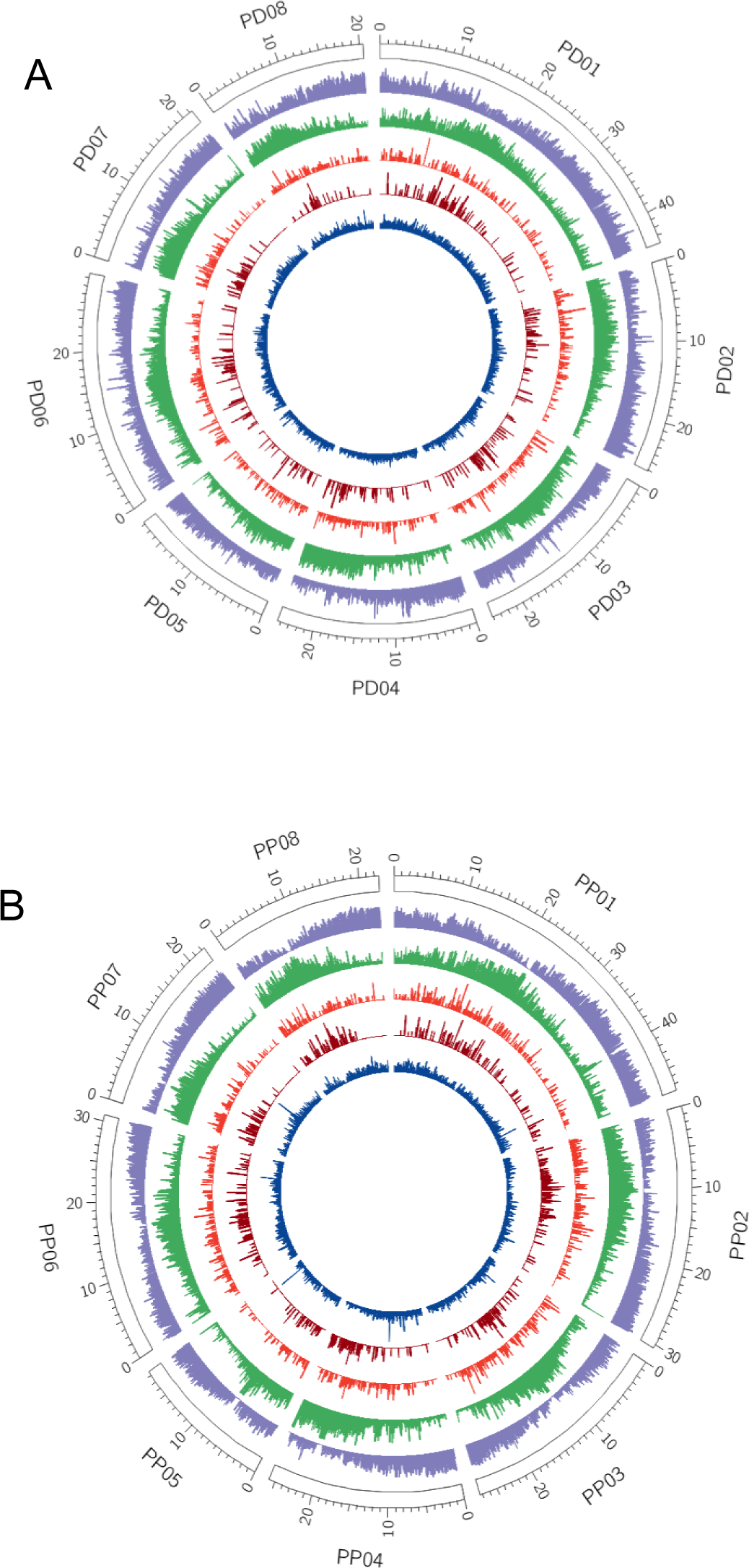
Distribution of gene and TE abundance along *P. dulcis* (A) and *P. persica* (B) chromosomes. Outer to inner tracks represent the coverage per 100 kb of Genes, TEs, Copia LTR retrotransposons, Gypsy LTR retrotransposons, and MITEs

In addition to the general TE annotation, we performed a dedicated annotation of the LTR-retrotransposons and MITEs in the almond and peach genomes. A conservative search for LTR-retrotransposons with a well-preserved structure (i.e. presence of LTRs and coding capacity for retrotransposon-related proteins) allowed annotating approximately 2,200 elements in both almond and peach (Table S7). Whenever possible, these elements were classified as Copia or Gypsy (i.e. when coding regions for both integrase and reverse transcriptase were detected, which allowed to classify them) or remained as unclassified LTR-retrotransposons. Although the content of these elements was similar in both genomes, the number of LTR-retrotransposons that remained unclassified in almond was slightly higher. The distribution of the almond and peach retrotransposons along chromosomes was also highly similar, with Gypsy elements showing a tendency to concentrate in a central region of chromosomes, probably coinciding with the centromeric regions, whereas Copia elements more evenly distributed (Figure 1). A conservative search for MITEs with well-preserved TIRs rendered 10,460 MITEs in almond and 8,738 MITEs in peach (Table S7). The distribution of these elements along chromosomes is similar in peach and almond and follows that of Copia LTR retrotransposons (Figure 1).

### LTR retrotransposon dynamics in almond and peach

To gain insight into the evolution of almond and peach LTR-retrotransposons, we grouped all almond and peach elements into clusters showing sequence identity higher than 80% along more than 80% of their length. Most of the elements (66.2%) were grouped into clusters of, at least, two elements. Two hundred and fifty-nine clusters (81%) were mixed-clusters, and contained 93% of the almond and peach elements. An analysis of the insertion times of these LTR-retrotransposons shows that the number of recent (≤5 Mya) LTR-retrotransposons is clearly higher in peach than in almond (Figure 2A). An analysis of the insertion time distribution of individual clusters within LTR retrotransposon families (belonging to Gypsy and Copia superfamilies, or that are unclassified), shows that many of them contain insertions that are younger in peach as compared with almond (Figure S7).

**Figure 2.**
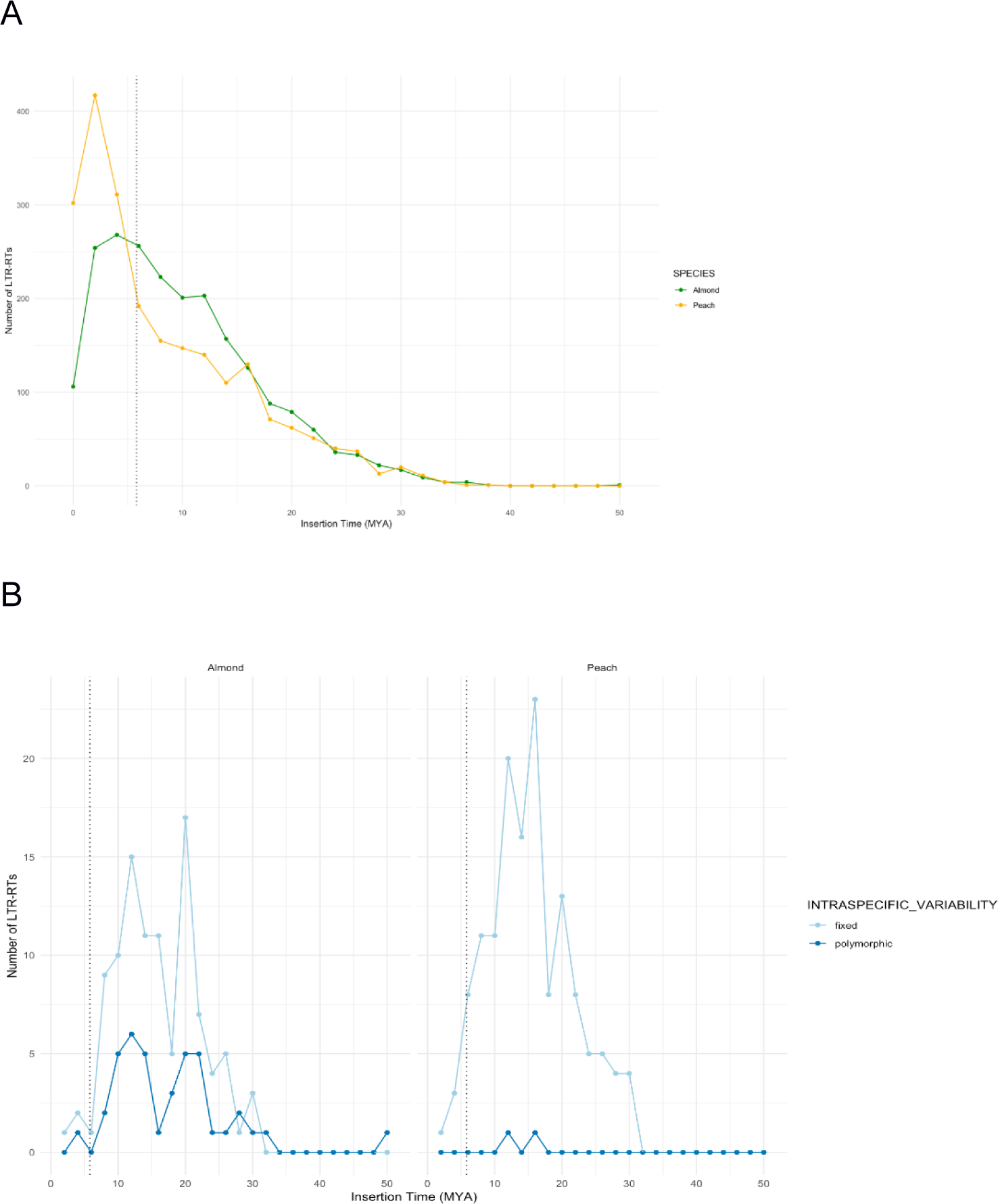
Dynamics of LTR-retrotransposons in peach and almond. **A.** Insertion time of complete LTR-retrotransposons in *P. persica* and *P. dulcis.* **B.** Insertion time of polymorphic and fixed orthologous LTR-retrotransposons in almond (left) and peach (right).

To further understand LTR-retrotransposon dynamics we analyzed the prevalence in the species of the LTR-retrotransposon insertions found in peach and almond reference genomes by analyzing resequencing data from ten peach and ten almond cultivars. This analysis shows that the LTR-retrotransposon insertions are frequently polymorphic among almond cultivars whereas they appear often fixed in peach. An analysis of the insertion time distribution of fixed and polymorphic LTR-retrotransposons in both species shows that, whereas an important number of LTR-retrotransposon insertions older than the estimated speciation time (see section 3.6) are polymorphic in almond, peach contains very few old polymorphic insertions suggesting that they were lost this species (Figure S8).

As all these analyses may be somewhat biased by the different quality of peach and almond genome assemblies, we performed a detailed analysis of the LTR-retrotransposon insertions comparing orthologous loci in both species. We were able to unambiguously identify the orthologous locus to 1,155 full-length LTR retrotransposon peach insertions and 1,134 almond insertions (around 51% of the insertions in both species). These correspond to 142 insertions found intact in both species (conserved insertions), 592 and 440 new insertions in peach and almond, respectively, 422 peach insertions partially deleted or rearranged in almond and 562 almond insertions partially deleted or rearranged in peach. An analysis of the ages of these LTR retrotransposon insertions belonging to the different categories showed that, as expected, the majority of the new insertions in both genomes were younger than the estimated speciation date. Almond contained a larger fraction of new insertions that are older, which probably corresponds to elements that were polymorphic in the ancestor and that were subsequently lost in peach. Also as expected, the vast majority of the conserved insertions were older than the estimated speciation time of peach and almond (Figure S9). In addition, while almost all of insertions were fixed in peach, an important fraction was polymorphic in almond (Figure 2B).

### Phylogenome analysis

To shed light on the evolutionary history of the genome of *P. dulcis* in the context of sixteen other sequenced plant species (Table S2), we generated the complete phylomes. These phylomes were filtered to remove the gene trees that contain proteins associated to transposons. After filtering, a total of 18,475 gene trees were kept for almond, and 20,812 gene trees, for peach. This filtered phylomes were scanned to infer duplications and speciation events and derive orthology and paralogy relationships from individual gene trees (Gabaldón, 2008).

To reconstruct evolutionary relationships, we concatenated the protein alignments of 262 genes, which had single-copy orthologs in all the 17 species considered. The resulting highly supported topology (Figure 3A) was congruent with current views on plant phylogeny (Shaw & Small, 2004) and results in *P. dulcis* and *P. persica* forming a clade, to the exclusion of *P. mume* and *P. avium* (Badenes & Parfitt, 1995; Scholz et al., 2013). The same topology was obtained when all individual gene trees were combined into a single species phylogeny by using a gene tree parsimony approach. We estimated the divergence times among these species using a Bayesian relaxed molecular clock approach. According to our results, *P. dulcis* diverged from *P. persica* approximately 5.88 Mya, from *P. mume* 20.84 Mya, and from *P. avium* 62.04 Mya (Figure 3A, Table S8).

**Figure 3.**
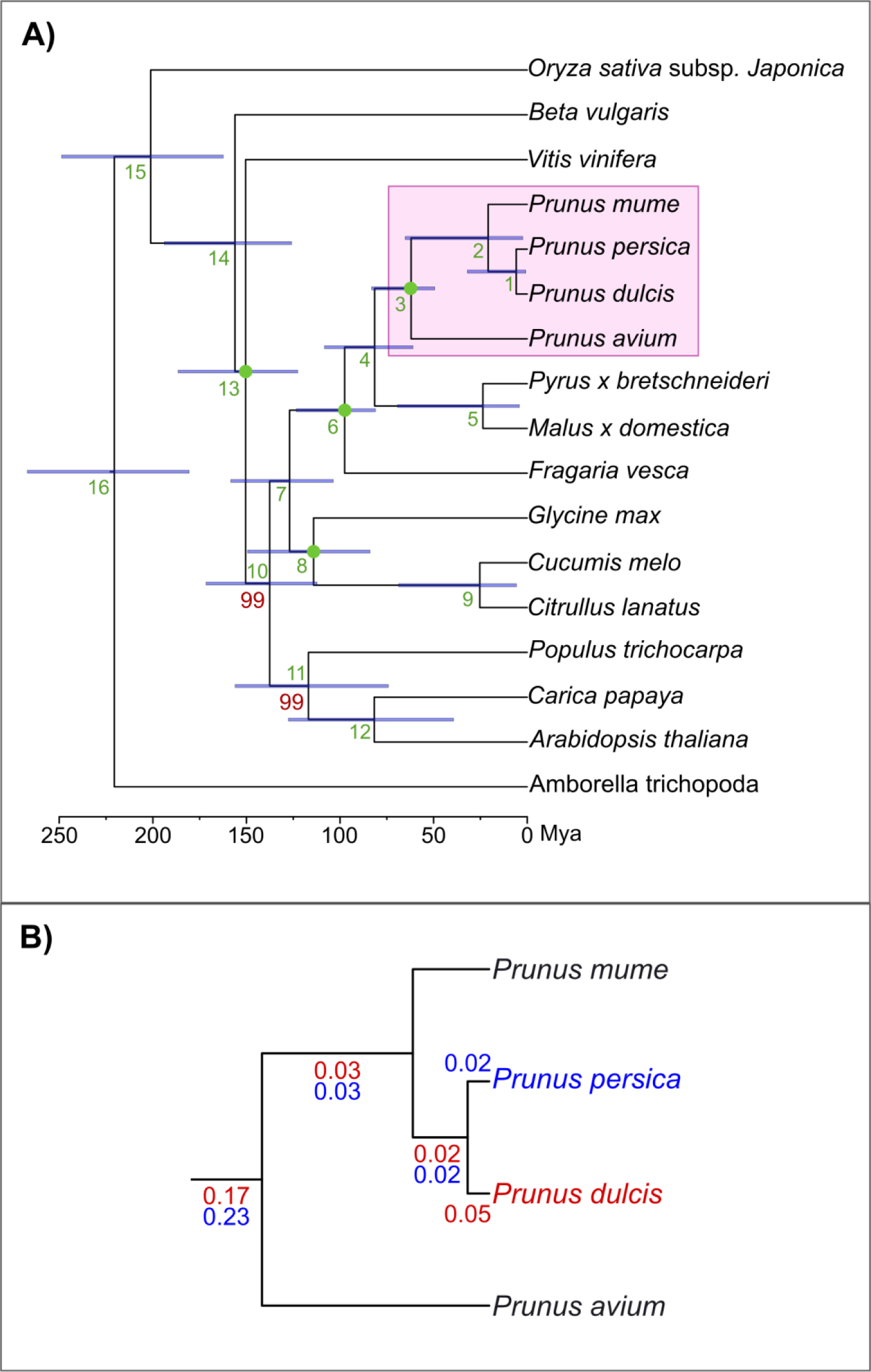
Species tree obtained from the concatenation of 262 widespread single-gene families. **A)** Full species tree. All *Prunus* species are highlighted in pink. All bootstrap values that are not maximal (bootstrap 100%) are indicated in red. Green numbers correspond to the nodes in Table S9. Bars at the nodes indicate the uncertainty around mean age estimates based on 95% credibility intervals. Scale at the bottom shows the divergence time in Mya. Green dots represent selected calibration points. **B)** Zoom in of the *Prunus* group. Numbers indicate the duplication ratio for each branch calculated with the phylome of almond (red) and peach (blue).

We then calculated the duplication frequency (i.e. average number of duplications per gene) for each node of the species tree, and observed a slightly high duplication frequency (∼0.20, Figure 3B) at the ancestral node of all *Prunus* species, which was supported by both almond and peach phylomes. A functional analysis of protein families duplicated at this node in each phylome shows enrichment of some molecular functions such as methyltransferase activity, ionotropic glutamate receptor activity, terpene synthase activity, oxidoreductase activity and transferase activity. In addition, some biological processes were enriched: response to auxin, metabolic process and oxidation-reduction process (Table S9).

Then we focused on duplications specific to each almond and peach, including large expansions. A total of 1,175 (4.4%) almond proteins and 831 (3.1%) peach proteins have an in-paralog (a recent paralog resulting from a duplication that specifically occurred in the almond and peach lineage, respectively). Those paralogs could be assigned to 542 almond specific gene expansions, and 367 peach specific gene expansions. In both, almond and peach, most expansions (540, 99.6% – almond; 363, 98.9% – peach) have a moderate size (2-5 in-paralogs, Figure S10). Some almond expansions of size two encode putative members of the lignin biosynthesis pathway (Vanholme et al., 2010) such as caffeic acid 3-O-methyltransferase (COMT, Prudul26A009858P1-Prudul26A011895P1), shikimate O-hydroxycinnamoyl transferase (HCT, Prudul26A003924P1-Prudul26A028947P1, Prudul26A000852P1-Prudul26A022843P2). Interestingly, these genes have undergone parallel duplications in *P. persica*, *P. mume*, and *P. avium*.

We next analyzed protein gains and losses in the lineages leading to almond and peach. When we analysed the almond phylome, a total of 1,471 proteins were gained in almond and 1,146 were lost in peach. For the peach phylome we found that 1,984 proteins were gained in peach and 3,157 were lost in almond. Functional analysis shows that the proteins lost in almond respect to peach are enriched in functions related to serine-type endopeptidase inhibitor activity, nutrient reservoir activity, lipid transport, response to auxin, oxidation-reduction process and ion transport. Conversely, genes lost in peach with respect to almond are mainly enriched in functions related to transferase activity, transcription and ATP synthesis coupled proton transport (Table S10).

### Variability of almond cultivars

#### Read mapping rate, depth and genome coverage

Alignment of reads from the ten almond varieties resulted in an average mapping rate of 89.2%. Specifically, 825,914,441 out of the 919,019,814 reads mapped onto the almond assembly, with Nonpareil and Vivot, and Falsa Barese and Genco showing the highest (94%) and lowest (79%) mapping rate, respectively (Table S11). Regarding sample depth and genome coverage, an average of 39.7 read depth was detected for the ten varieties whereas 93% of the assembly was covered by the resequencing data on average. Marcona and Falsa Barese had the highest (51.4X) and lowest (27,7X) read depth, respectively, whereas Nonpareil and Falsa Barese had the highest (95%) and lowest (90%) coverage, respectively (Table S11).

#### Variant calling and phylogenetic analysis

Genetic variabiltity analysis resulted in the detection of 2,253,377 variants composed of 2,203,582 (87%) SNPs and 330,795 (13%) INDELs. Genome-wide distribution of SNPs and INDELs can be seen in Table S12. Nonpareil had the highest number of SNPs and INDELs, 1,072,759 and 142,142, respectively, whereas Ripon had the lowest number of SNPs and INDELs, 827,397 and 94,070, respectively. Average SNP density was calculated as 6.2 SNPs/Kb, whereas the average heterozygosity for the ten cultivars was 0.44 (Table S13). Our results on SNP density and heterozygosity for almond are lower (SNP density of 19.1 and heterozygosity of 0.69) from those published recently (Yu et al. 2018) most probably due to the use of the peach genome as a reference sequence for the variant calling, a varietal panel of larger genetic diversity, or different filters and tools used for variant calling, which overall may have resulted to the additional detection of homozygous variants in the almond varieties.

A graphical representation of SNP and INDEL distribution in windows of 100 kbp showed a similar profile for most of the almond varieties analyzed (Figure S11). Nevertheless, Ripon and Falsa Barese displayed lower variant density in the region from 28Mb to the end of Pd01 and in the region from 13Mb to the end of Pd07. Also local decrease in the number of SNPs and INDELs could be detected for specific varieties by examining the distribution profiles (Figure S11).

The analysis of deletions in the collection of almond cultivars is presented in Table S14. For small deletions (1-50 bp), numbers were about half of those estimated for INDELs in Table S12 (∼60,000 vs ∼120,000 per cultivar), as expected considering that only deletions and not insertions are taken into account. On average, 27% of these deletions overlapped with TEs. Considering large deletions (>50bp), we detected 1,219 unique events, the majority of which fell within the 51-500bp range, with an average number of 88 deletions per variety. Marcona had the largest number of deletions (219) whereas Ripon had the lowest number (12) (Table S14). Five hundred and eighty-eight large deletions (48.2% of the total) overlapped with TE elements. For deletions larger than 500bp, almost all the events were found to overlap with TEs (92.6% at range 501-10000 and 100% at range 10001-50000; Table S14; Figure S12). Overlap with TEs was very similar among varieties and represented an average of 48.2% of large deletions, with Nonpareil and Ripon showing more than 50% of overlap with TEs and Cristomorto with the least percentage of overlap (38.8%).

A SNP-based phylogenetic analysis grouped the almond varieties into two main clades where the first clade contained Cristomorto, Falsa Barese and Genco whereas the second clade was split into two subclades; the first containing Aï, Belle d’Aurons, Nonpareil and Ripon and the second Desmayo largueta, Marcona and Vivot (Figure S13). These data of genetic relatedness are in agreement with the geographical origin of the analyzed cultivars, grouping the Italian cultivars, the French and US cultivars and finally the Spanish cultivars in the same clade. The fact that French and US cultivars are clustered together is in agreement with the known origin of US materials coming from French imported accessions (Kester et al. 1991).

#### Analysis of indel variants between peach and almond genomes and its relationship with TE sequences

To assess the structural variability between almond and peach genomes we aligned almond genome contigs to the peach reference genome. A total of 92.96% of the almond Texas assembly could be aligned with that of the reference peach genome sequence of Lovell with an average identity of 95.59%. We detected a total of 20,418 variants accounting for 18 Mb of sequence, equivalent to 8% of the almond genome (Table S15). Whereas only a 25% of indels between 50-500 bp colocalized with annotated TE insertions, 65% of indels in the range of 500-10,000 bp and the vast majority (86%) of indels in repetitive regions involved TEs.

Similarly to the analysis of deletions reported in the previous section with almond cultivars, we mapped resequencing data from the peach variety Earlygold with the almond reference genome. The number of deletions found in an average almond cultivar compared with the almond reference sequence was on average 62,238, whereas that of the peach cultivar studied was more than double (126,137) (Table S14). However, when taking only those of larger size (>50 bp), peach deletions were more than tenfold (1,436 vs. 120) more frequent than almond deletions.

### Transposon-induced variability in peach and almond traits

The almond fruit resembles that of peach and other *Prunus* species, the major differences being that in almond the mesocarp does not develop to produce the fleshy tissue typical of some other *Prunus* fruits, and that the almond seed does not accumulate the high levels of the cyanogenic diglucoside amygdalin that renders the seeds of peach and other *Prunus* bitter and toxic. In order to shed light on the genetic differences underlying these phenotypic differences we have compared the genomic regions containing the genes known to control the expression of these characters in both species.

It has been recently shown that the sweet almond phenotype is due to the reduced expression of the genes encoding two cytochrome P450 enzymes catalyzing the first steps of the amygdalin biosynthesis in sweet almond varieties as compared with bitter almond varieties (Thodberg et al. 2018). It has also been shown that this reduced expression is not related with differences in the gene sequence, which points to a difference in the regulation of the expression of those genes (Thodberg et al. 2018). We have compared these two almond genes with their homologs in peach, sweet cherry and *P. mume* and found that they are highly conserved (not shown). However, one of the two genes, *CYP71AN24*, is flanked by several almond-specific TE insertions, and in particular two MITE insertions in its proximal upstream region (Figure 4A). A preliminary analysis of the methylation of this region shows that the almond-specific TE insertions flanking the *CYP71AN24* gene are highly methylated. The insertion of TEs in the proximal upstream region of the *CYP71AN24* gene may have affected directly or indirectly its expression due to its high methylation. In addition, the presence of the TEs also correlates with a much higher methylation of this gene in almond as compared with peach (11.6% methylation at CG, and 0,1% methylation at CHG in almond versus 0,1% methylation at CG, and 0,02% at CHG in peach), which may be the result of methylation spreading from the TEs. An analysis of the structure of this locus in *P. webbii*, a wild species closely related to almond which produces bitter seeds, and in two almond sweet varieties (Cristomorto and Marcona) and two almond varieties producing bitter seeds (D05-187 and S3067), shows that the presence of the TE insertions (and in particular that of the MITE named TIR2), perfectly correlate with the sweet vs. bitter seed phenotype (Figure 4B).

**Figure 4.**
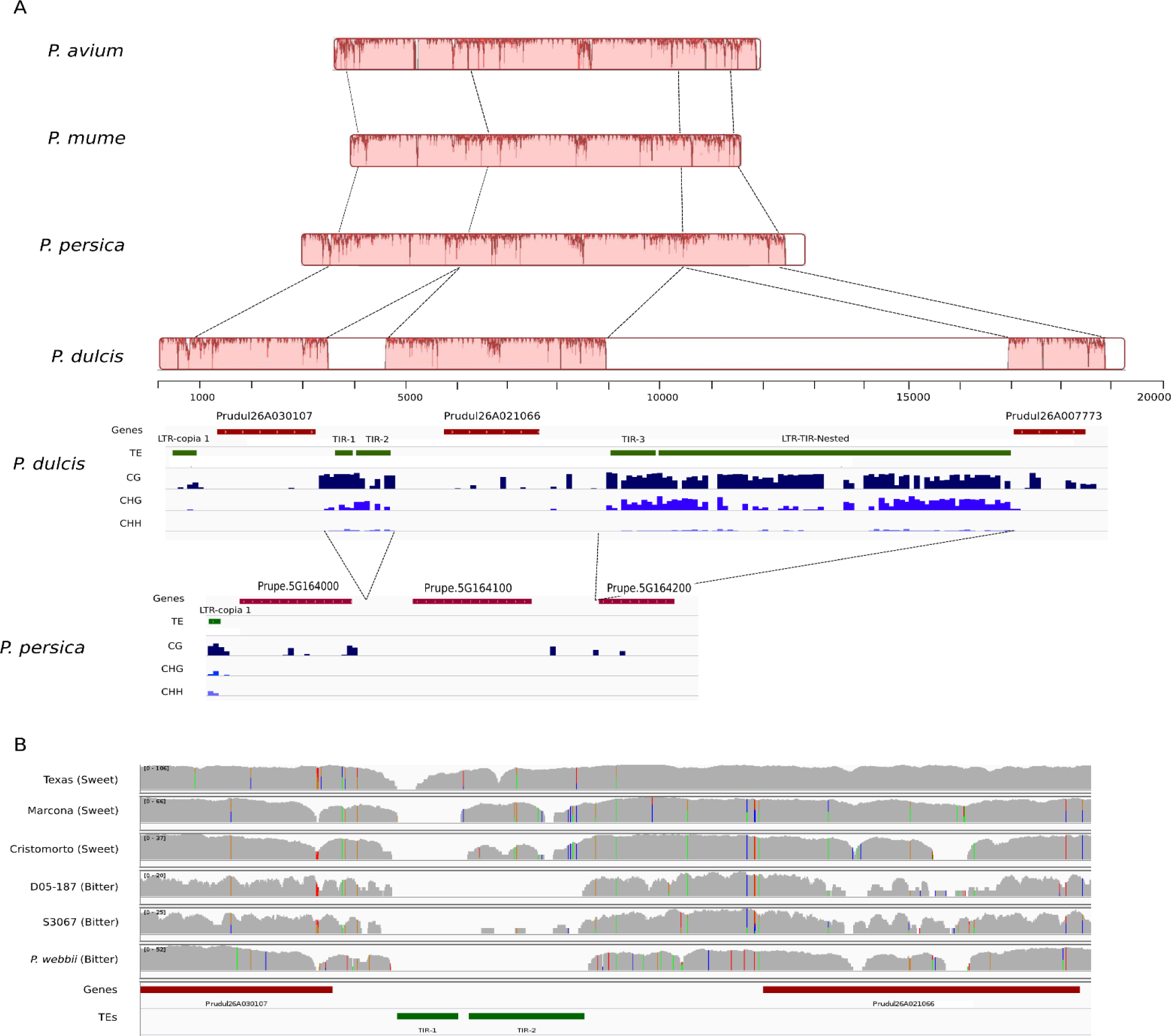
Analysis of the locus of the CYP71AN24 gene in almond varieties and *Prunus* related species. A. Nucleotide conservation of the CYP71AN24 region between *P. avium, P. mume, P. persica* and *P. dulcis* based on a Mauve multiple alignment (physical distance scale is in bp). White boxes represent inserted regions in *P. dulcis*. IGV tracks of the gene and TE annotations of *P. dulcis* and *P. persica* along with their DNA methylation levels in the three different contexts (CG, CHG and CHH) are shown below. B. IGV spnapshot of the region containing the CYP71AN24 gene and the polymorphic TE insertions displaying the coverages of mapped DNAseq reads from resequencing data of sweet- and bitter-kernel *P. dulcis* varieties, as well as from that of the closely related *P. webbi*.

The analysis of model plant species, such as Arabidopsis and tomato, has shown that the development of the fruit is the result of the combined regulation of meristem organization, flower development, and fruit cell proliferation and expansion. We selected 97 genes (Table S16) that belong to gene families involved in these processes, including WUSCHEL (WUS) and CLAVATA (CLV) genes involved in the control of the meristem and whose mutants lead to larger fruits in tomato and genes known to control fruit shape (Rodriguez et al. 2011). We have analyzed the structure of these genes in peach and almond and found that six of them present differential TE insertions within the genes in their proximal (less than 1000 nt) upstream region that probably contains their promoter. These species-specific TE insertions are all highly methylated and may have altered their expression regulation (Table S17). Moreover, and in addition to the potential mutation of transcriptional regulatory elements, MITE insertions could have provided new transcription factor binding sites (TFBS), as MITEs have been shown to frequently amplify and mobilize TFBS in plants (Morata et al 2018).

## Discussion

Using a hybrid strategy based on short and long read DNA sequences and the information provided by existing linkage maps we have assembled the highly heterozygous genome of almond cv. Texas into a highly-complete, contiguous and low-redundant assembly with eight pseudomolecules corresponding to the eight chromosomes and comprising 227.6 Mbp of sequence. Annotation of this genome has resulted in the identification of 27,042 protein-coding genes and 6,800 non-coding transcripts. The assembled sequence was highly syntenic with the genome sequence of peach (Verde et al. 2013) as expected considering previous information on the close genetic similarity between these two species (Dirlewanger et al. 2004).

Based on molecular data and the use of fossil records we estimated the divergence times of *P. dulcis* with respect to other sequenced *Prunus* and a diverse sample of 17 plant species. Our estimate of 5.88 Mya for the divergence of peach and almond from a common ancestor is similar to that of recent molecular evidence that places this figure between 5.0 and 8.0 Mya (Delplancke et al. 2013; Velasco et al. 2016; Yu et al. 2018). This is compatible with the separation of the ancestral species by the uplift of the Central Asian massif in two subpopulations that faced completely different environments: one (almond and its close relatives) in the arid steppes of central and western Asia and the other (peach) in the subtropical climate of southwestern China, near to where the first fossil endocarps were found dated of 2.6 Mya (Su et al. 2013).

In agreement with the results of other sequenced diploid *Prunus* genomes (Verde et al. 2013; Zhang et al. 2012; Shirasawa et al. 2017; Baek et al. 2018) we have not found evidence on any recent whole-genome duplication of almond. Analysis of duplicated gene sequences indicated a parallel gene expansion for all sequenced *Prunus* species genomes for genes involved in lignin biosynthesis, such as caffeic acid 3-O-methyltransferase and shikimate O-hydroxycinnamoyl transferase. One of the distinctive aspects of *Prunus* is that its fruit is a drupe, characterized by the formation of a strongly lignified mesocarp (the “stone”), unlike most of its closest taxa that have follicetum and nuculanium as fruit types (Xiang et al. 2016). These duplications may be at the origin of the formation of the stone, determine its characteristics and be crucial to understand its evolution and possible modification, with important consequences for plant breeding, including the production of stoneless cultivars in *Prunus* fruit (Callahan et al. 2015).

As we have shown from the comparison of the peach and almond reference genomes, as well as the analysis of the structural polymorphisms between both genomes, insertion/deletion events seem to explain a substantial part of the divergence of peach and almond genomes from their common ancestor. In this paper we show that most of such structural differences, particularly those of larger sizes, were related to TE sequences. LTR-retrotransposons and MITEs constitute the two most prevalent superfamilies of TEs in plants (Casacuberta and Santiago 2003). While the proportion of the almond genome consisting of TEs (38%) was similar to that of other sequenced genomes of similar size (i.e. from 30% in peach (Verde et al. 2013) and 43-47% in other *Prunus* (Zhang et al. 2012; Shirasawa et al. 2017; Baek et al. 2018) and with a very similar distribution of TEs along the almond and peach chromosomes, the detailed analysis of the dynamics of LTR retrotransposon evolution revealed key aspects of the divergence of almond and peach genomes after their speciation. In short, almond has maintained more ancestral LTR-retrotransposons, which are still in some cases polymorphic within the species. Conversely, peach has lost most polymorphic ancestral insertions but has undergone a higher recent retrotransposition activity. Slight differences in the mechanisms controlling transposon activity, triggered in peach by strong genetic changes determined by the adoption of the self-compatible mating behavior (Yu et al. 2018), could explain a higher LTR-retrotransposon activity in this species. Additionally, the contrasting data on the polymorphism of ancestral TEs between peach and almond may be explained by i) the mating types: selfing in peach and outcrossing in almond, and ii) differences in their recent history, with a strong reduction of population size in peach prior to its recent expansion as a cultivated species 2,000 years ago (Velasco et al. 2016; Yu et al. 2018), while almond population sizes remained higher (Yu et al. 2018). Further analyses should help to clarify this in the future. In any case, the results presented here suggest that LTR-retrotransposons, and in general TEs, may explain an important fraction of the interspecific variability between peach and almond, as well as the intraspecific variability of both species.

LTR-retrotransposons are at the origin of somatic mutations in plant species, some of which of high agricultural value (Foster and Aranzana, 2018). Only in peach, white vs. yellow fruit color (Falchi et al, 2013) hairy vs. glabrous fruit (Vendramin et al, 2014) and stonyhard vs. melting flesh texture (Tatsuki et al. 2018) are caused by the action of transposon movement. MITE insertions have also been linked to crop traits, such as sex determination in melon (Martin et al. 2009) or a drought tolerance phenotype in maize (Mao et al. 2015). The analysis of almond × peach interspecific progenies identified 11 Mendelian genes explaining the inheritance of some key agronomic characters: one of them, responsible for the formation of a thick mesocarp that constitutes the peach flesh, and another, conferring juiciness to the fleshy mesocarp, both critical aspects of the difference between almond and peach fruits (Donoso et al. 2016). Our results suggest that TEs could be responsible for some of the genomic changes at the origin of the agronomic traits that distinguish peach from almond, such as mesocarp development and bitterness of the kernel. For one of them, sweet vs. bitter kernel, essential for the domestication of the almond, we show here that the sweet almond phenotype correlates with TE insertions surrounding gene *CYP71AN24*, involved in the synthesis of one of the key enzymes of the amygdalin pathway (cytochrome P450), and that could be responsible for the lower expression of this gene in the seed tegument of sweet almonds (Thodberg et al. 2018). Additionally, we identified six more genes related with fruit flesh formation that had similar patterns differing between almond and peach. We have not been able to associate any of these genes with the position of major genes or QTLs described so far in these species, although they are clear candidates for further studies.

The genome sequence of almond that we present in this paper will foster the knowledge on the genetics and breeding of this species by providing useful information on genes and markers with an unprecedented level of detail. For example, in addition to the variability for SNP markers that is described here, we found a high number of insertion/deletion variants of sizes >20 within almond and many more between almond and peach, that can be used as simple and cheap markers for genetic analysis as it has been done in other species (Wu et al. 2013; Li et al. 2015; Jain et al. 2019). *Prunus* crops are unique in the sense of being an economically important group of species that share an essentially common genome (Aranzana et al. 2019). Knowledge of the genome sequence opens new venues for the understanding of the evolution of closely related species that have in addition a slower rate of evolution due to their long intergeneration period, which may allow to detect modes and aspects of evolution that could be different or difficult to identify in herbaceous crops.

## Acknowledgments

This research was supported in part by grants from: the Ministry of Economy and Competitiveness (MINECO/FEDER projects AGL2012-40228-C02-01, AGL2015-68329-R and RTA2015-00050-00-00, Severo Ochoa Program for Centres of Excellence in R&D 201-2019 SEV-2015-0533 and CERCA Programme-Generalitat de Catalunya) from Spain. MW acknowledges grant support from the Waite Research Institute of the University of Adelaide.

## Author contributions

PA, TA, JMC and TG designed the project and supervised research. TA, FC, LF, JGG, MG, PR and AV performed the genome assembly and annotation. JMC, AB, FB, RC, JM provided the analysis of transposable elements. TG and IJ performed the phylome analysis. KA performed the variability and synteny analysis. AD, HD, AFM, MJRC, MW and WH collected materials, extracted DNA and provided resequencing data. JLG and BG generated the fosmid library. PP and TA provided long range DNA sequencing. PA, TA, JMC, JGM, TG and KA wrote the paper.

## Supplementary Material

Tables S1 to S17

Figures S1 to S13

