## Supplementary material for "Transposons played a major role in the diversification between the closely related almond (*Prunus dulcis*) and peach (*P. persica*) genomes: Results from the almond genome sequence": Suppl files

### Supplementary Figures

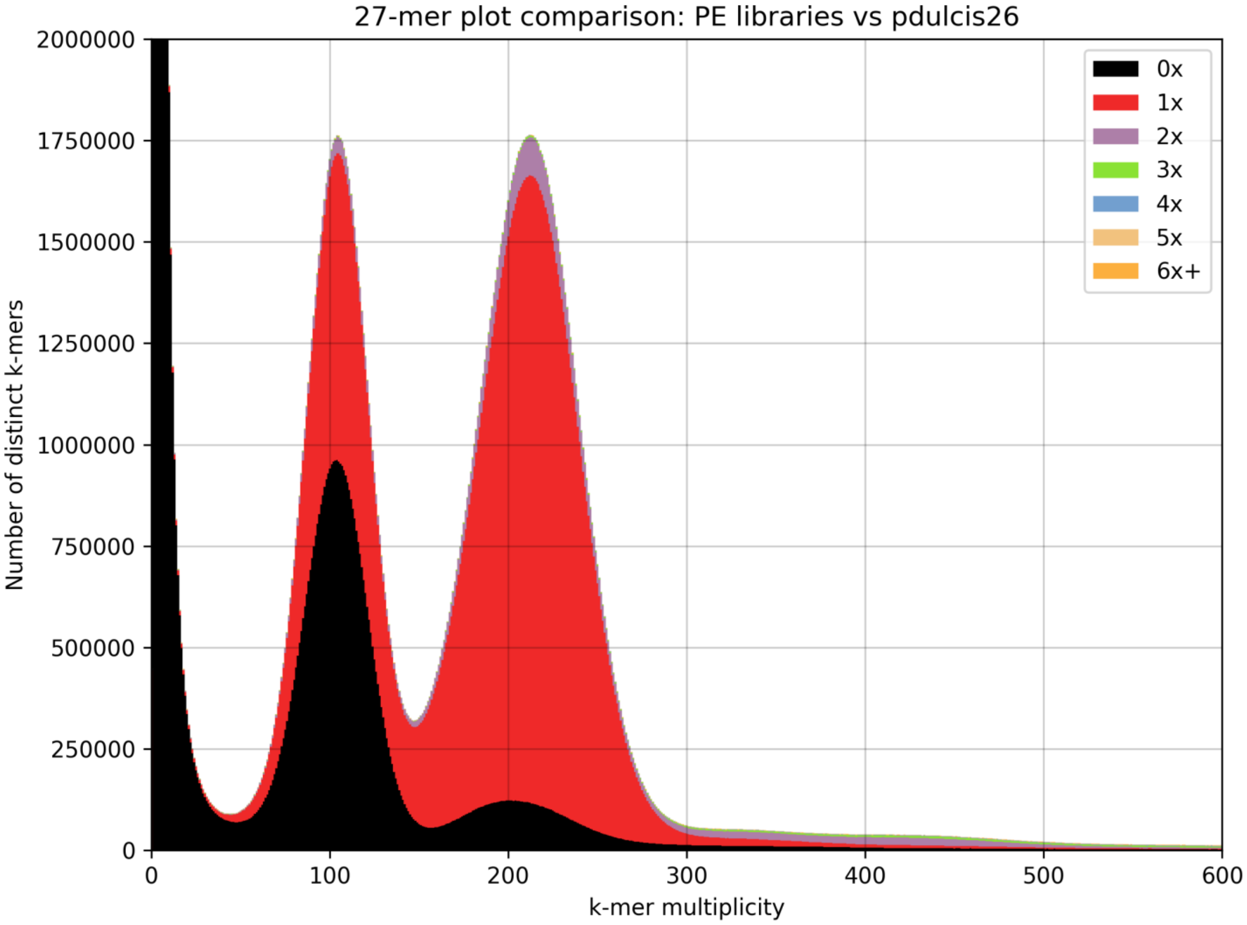

**Figure S1**. A stacked histogram based on the 27-mer matrix of the assembly and the paired-end Illumina libraries. The y-axis showing the number of distinct 27-mers in *Prunus dulcis* 2.0 (pdulcis26) and the x-axis the sequencing depth of each 27-mer in the PE libraries. The plot shows the amount of distinct K-mers absent from the assembly (0x class, in black), present in one copy (red), two copies (purple), and so on. As expected from a well-assembled diploid genome, half of the k-mers in the heterozygous peak (~105x) have been collapsed (the assembly contain just one allele) and the vast majority of the homozygous k-mers are present, with some missing (black) and some duplicated (purple).

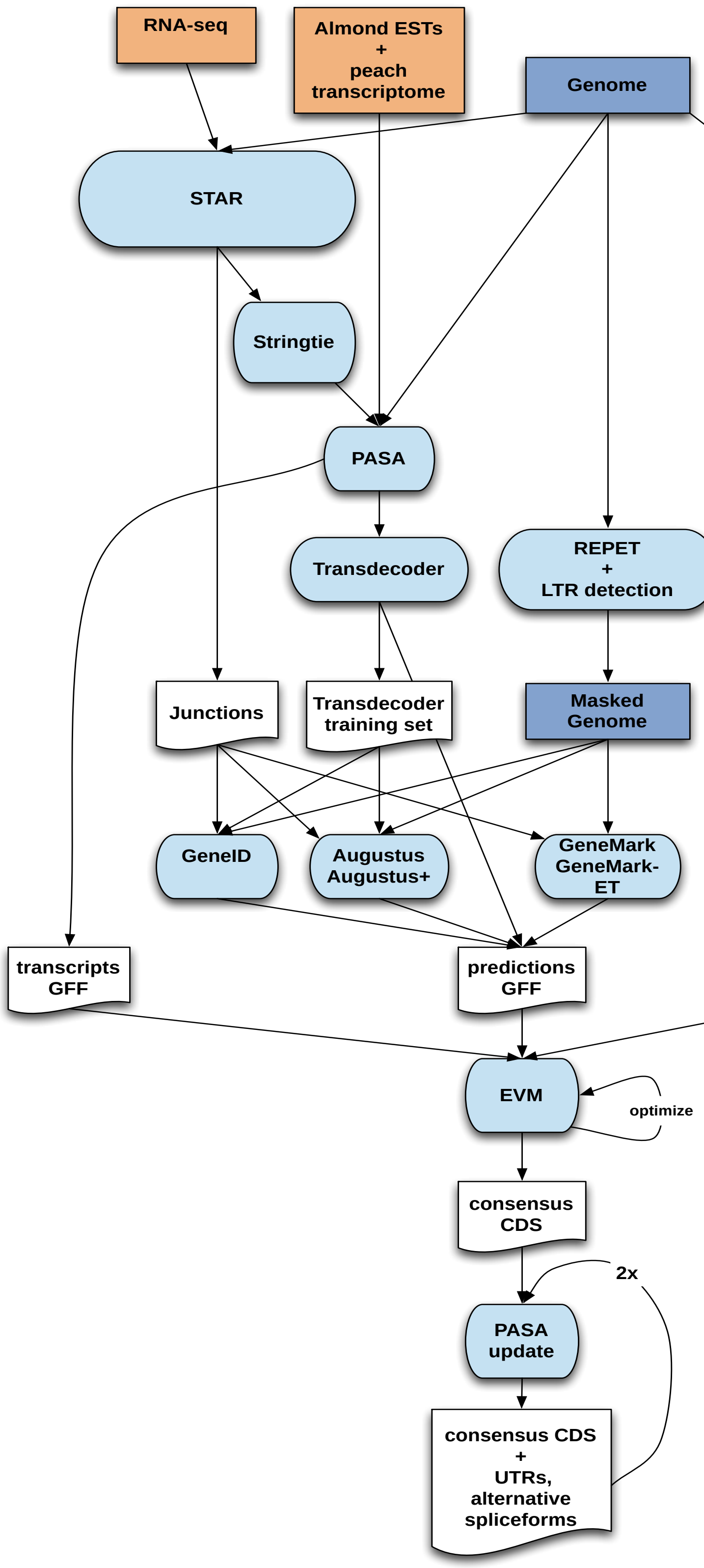

**Figure S2**. Protein-coding gene annotation pipeline.

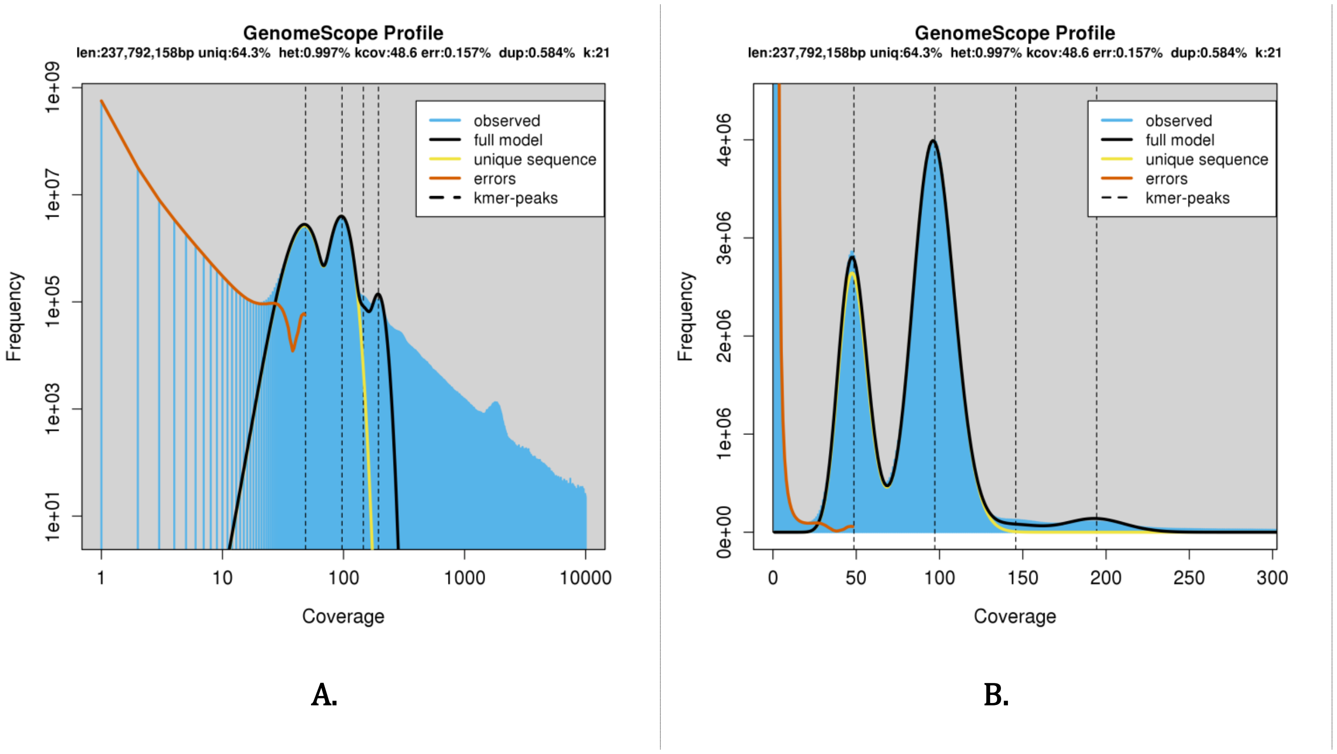

**Figure S3.** Genomescope k-mer coverage model fit using 21-mers. The 317 bp-insert 2x100 Illumina PE library was used for estimation. A) Both axes are shown in log-scale. The main homozygous peak is found at coverage 99, while the heterozygous peak is located at coverage 48.6. Two- and three-copy repeats are visible as well as a higher coverage peak corresponding to relatively few distinct k-mers that likely represents the chloroplast genome. B) A linear-scale plot focusing on the main and heterozygous peaks.

**Figure S4**. Synteny between the almond genome and the TxE linkage map

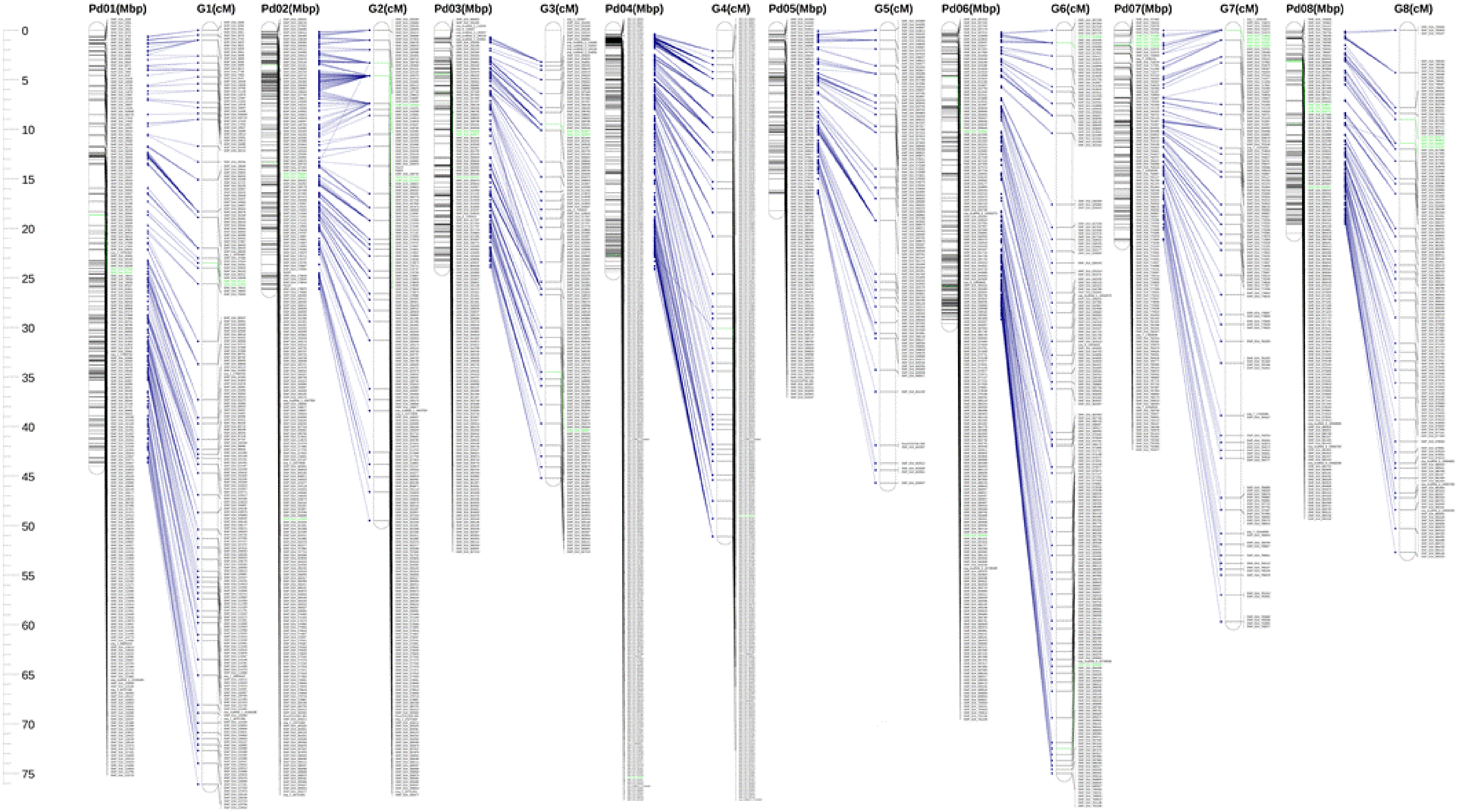

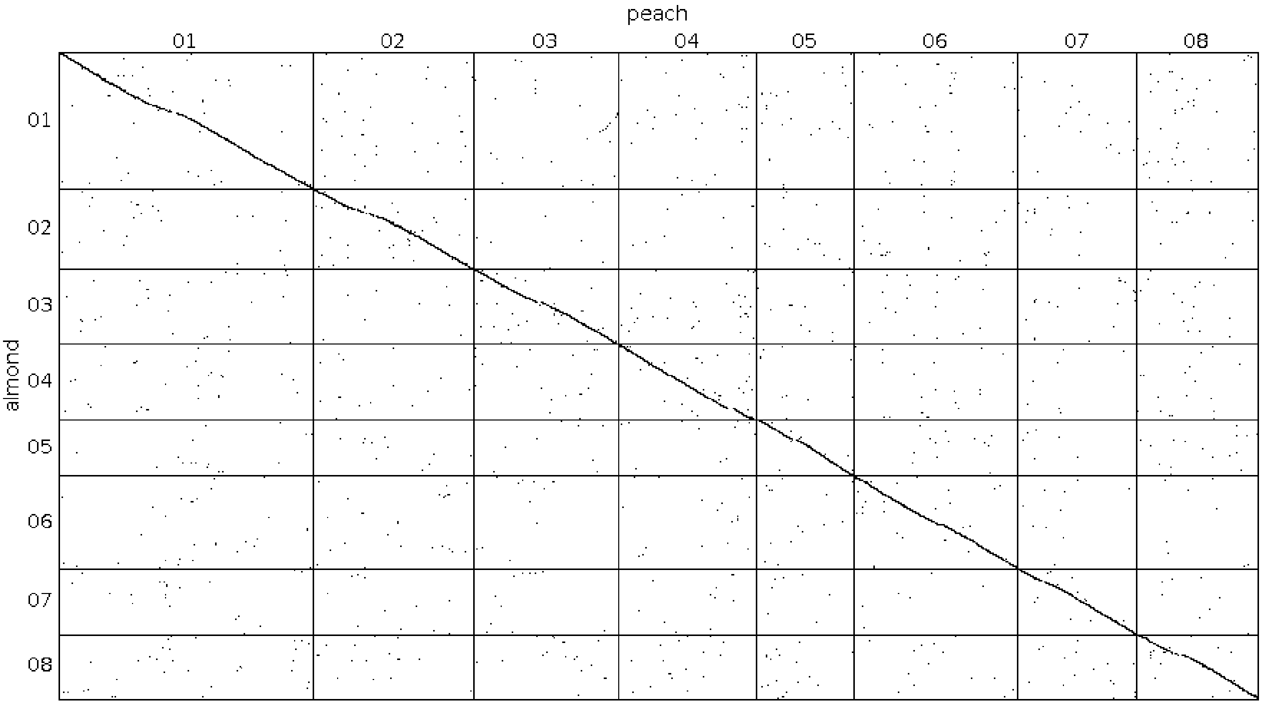

**Figure S5.** Synteny analysis of almond versus peach genomes.

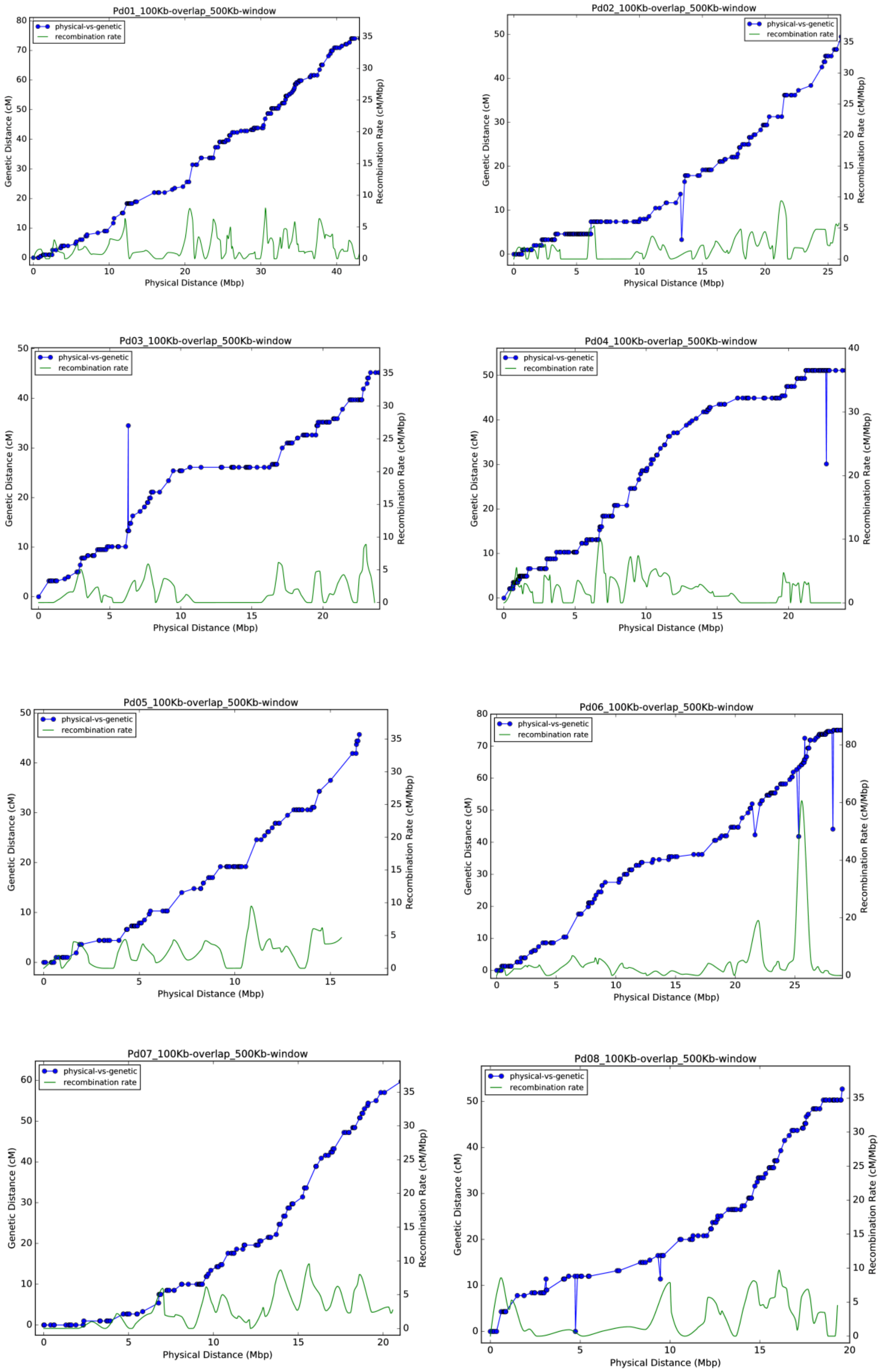

**Figure S6.** Distribution of recombination along chromosomes in almond. Recombination rates were calculated for 100 kb – overlapping windows with a size of 500 kb.

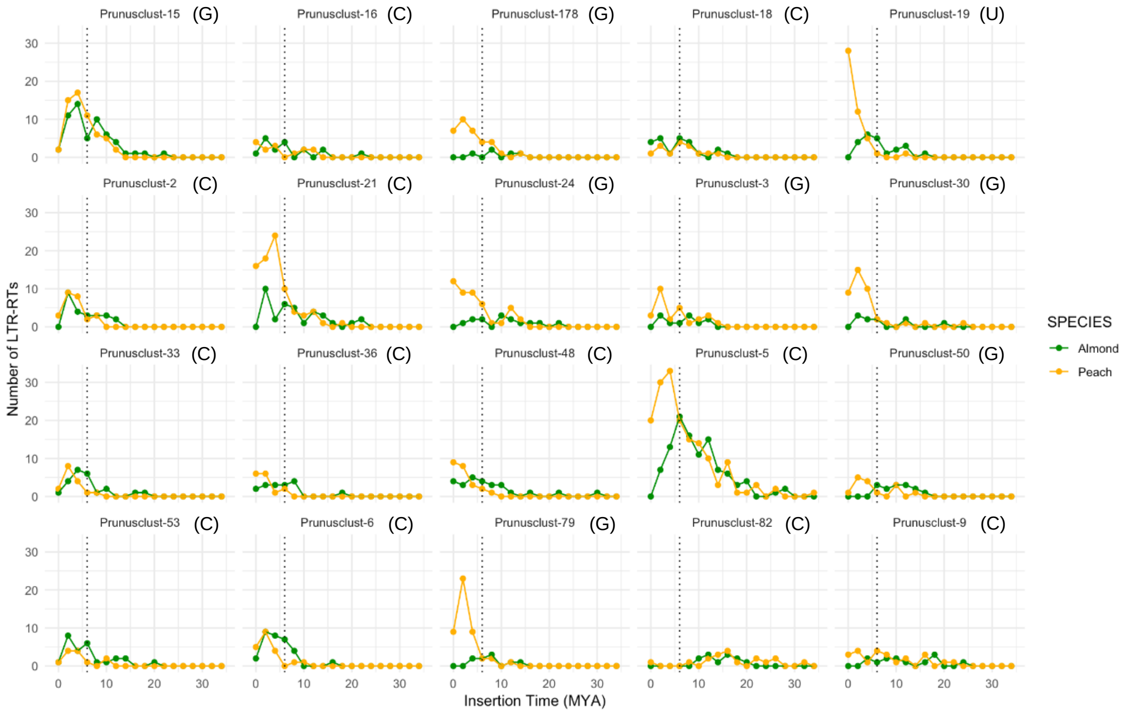

**Figure S7.** Insertion time distribution of individual LTR-retrotransposon families of the Copia (C) and Gypsy (G) superfamilies or that remained unclassified (U)

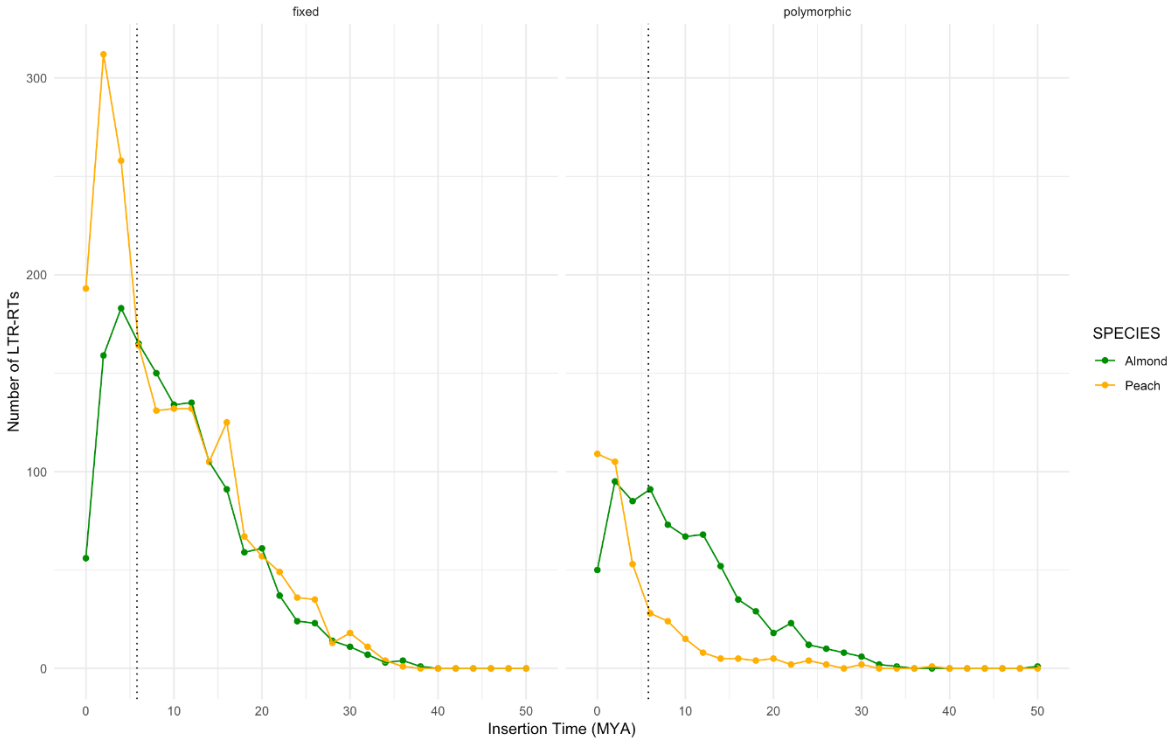

**Figure S8.** Insertion time distribution of fixed (left) and polymorphic (right) LTR-retrotransposon insertions in peach and almond

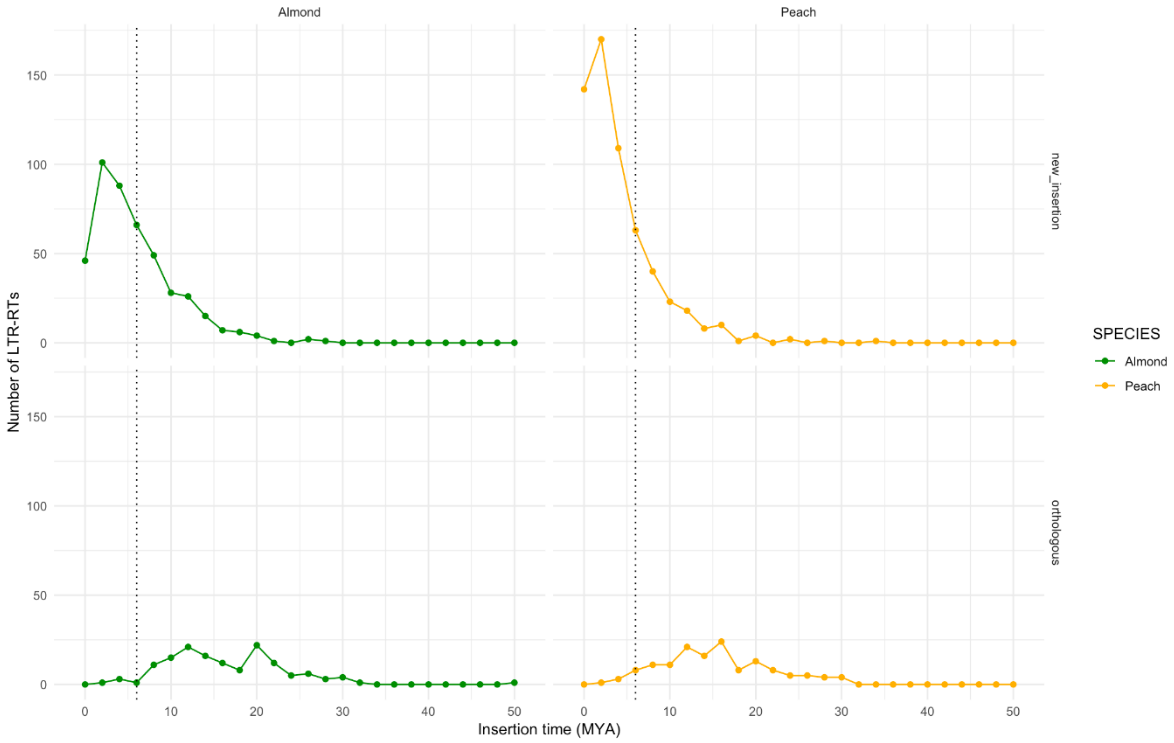

**Figure S9.** Insertion time distribution of new (upper panels) and orthologous (bottom panels) LTR-retrotransposon insertions in peach (left) and almond (right)

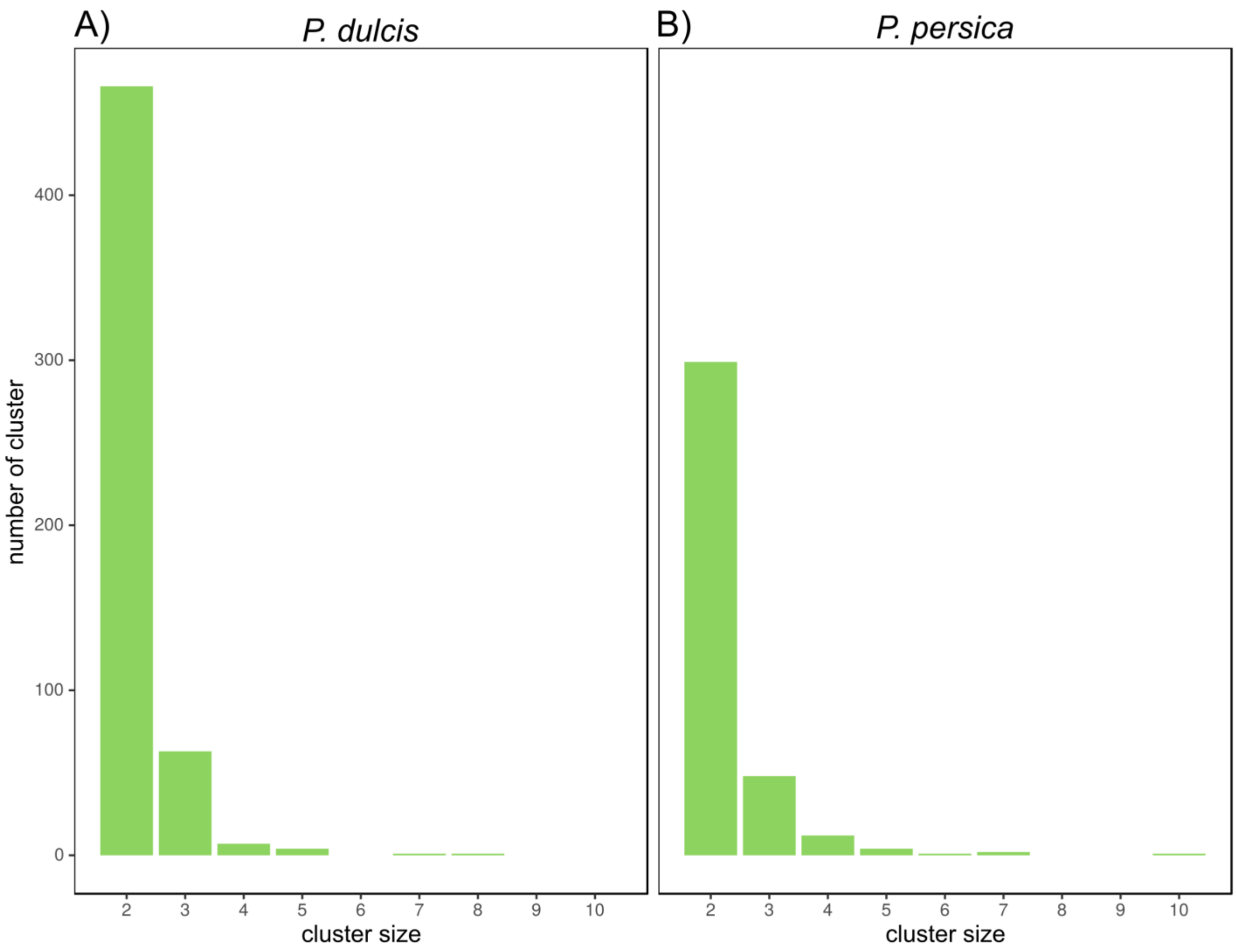

**Figure S10. Distribution of size of in-paralog groups resulting from species-specific duplications. A)** Almond-specific expansions. **B)** Peach-specific expansions.

(A)

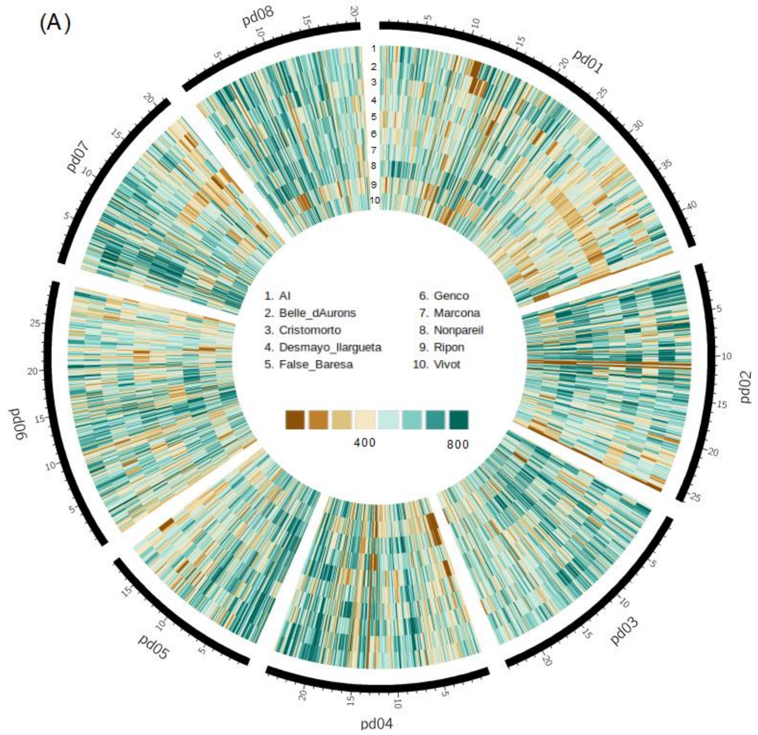
pd01
pd02
pd03
pd04
pd05
pd06
pd07
pd08
1. Al
2. Belle_dAurons
3. Cristomorto
4. Desmayo_llargueta
5. False_Baresa
6. Genco
7. Marcona
8. Nonpareil
9. Ripon
10. Vivot
400
800

(B)

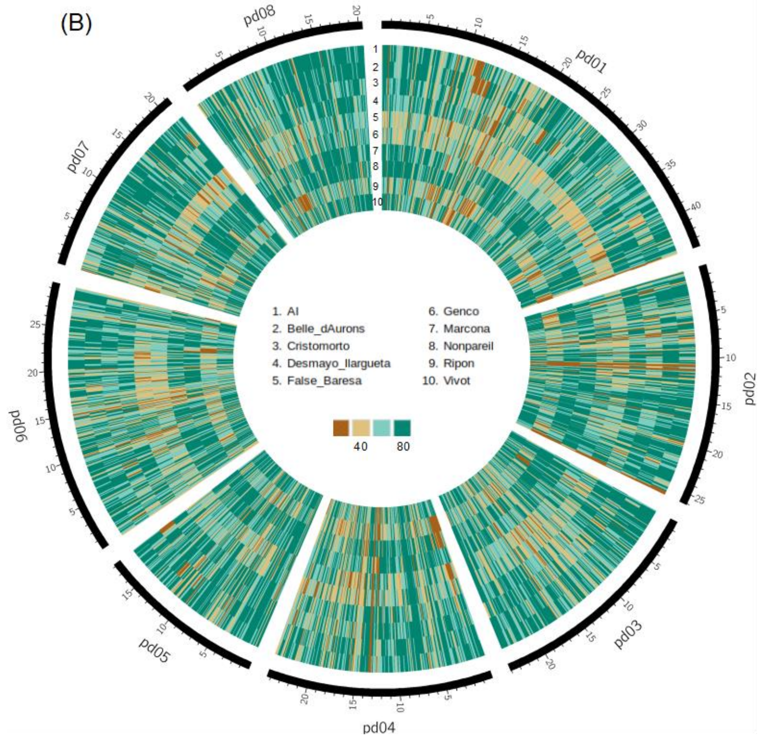
pd01
pd02
pd03
pd04
pd05
pd06
pd07
pd08
1. Al
2. Belle_dAurons
3. Cristomorto
4. Desmayo_llargueta
5. False_Baresa
6. Genco
7. Marcona
8. Nonpareil
9. Ripon
10. Vivot
40
80

**Figure S11.** Circos graphical representation of SNP (A) and INDEL (B) distribution across the almond genome. SNPs and INDELS from the ten resequenced varieties were binned in windows of 100Kb and their number per window was graphically plotted.

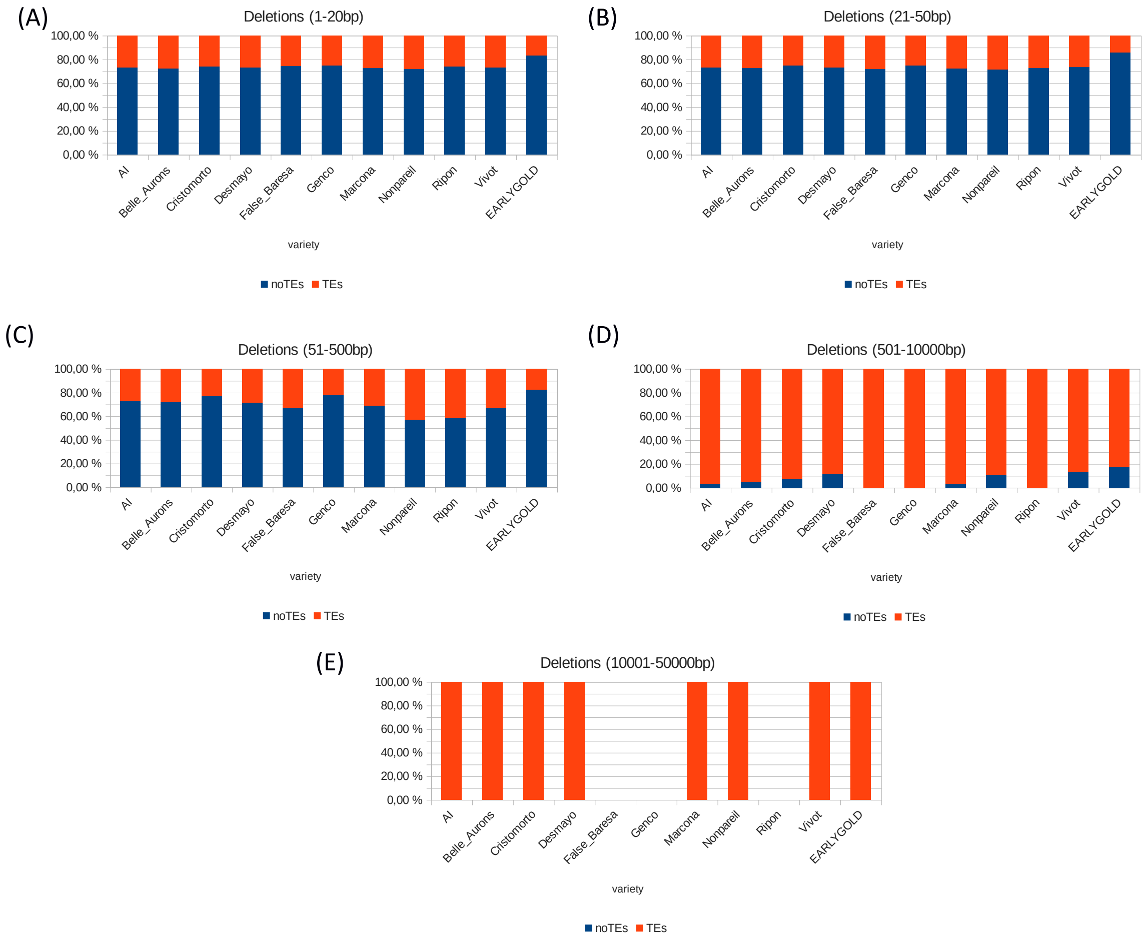

**Figure S12.** Percentages of non-TE and TE events for the different deletions in 10 almond varieties and 1 peach variety. (A) 1-20bp, (B) 21-50bp, (C) 51-500bp, (D) 501-10000bp and (E) 10001-50000bp.

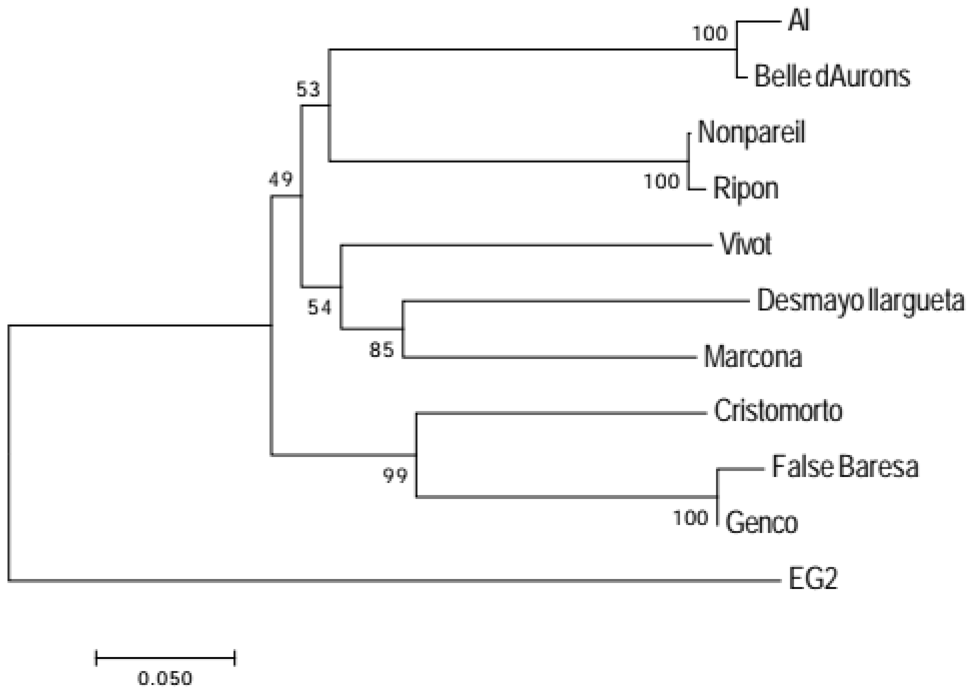

**Figure S13.** SNP-based phylogenetic analysis of the almond varieties.

#### Supplementary Tables

**Table S1.** Summary of sequence data used for Texas almond genome sequencing

| type | PE | PE | PE | MP | MP | Fosmid pool (mean) | ONT |
|---|---|---|---|---|---|---|---|
| **Fragment size** | 263 | 317 | 354 | 3.1kb | 5.2kb | 310 | HMW |
| **Read length** | 2 x 101 | 2 x 101 | 2 x 101 | 2 x 101 | 2 x 101 | 2 x 151/251 | N50=7.3kb |
| **Yield (Gb)** | 22.4 | 45.7 | 32.7 | 16.4 | 17.0 | 5.2 | 10.2 |
| **Depth** | 81 | 166 | 119 | 60 | 62 | >200 | 37 |

**Table S2.** List of species used in the phylome reconstruction. First column indicates taxa id, second column contains the species name and the third shows the source for the protein and the coding DNA sequences.

| TaxID | Species name | Source of protein coding sequences |
|---|---|---|
| FRAVE | *Fragaria vesca subsp. vesca* | JGI |
| PRUMU | *Prunus mume* | NCBI |
| AMBTC | *Amborella trichopoda* | Uniprot |
| BETVU | *Beta vulgaris* | CRG Ultrasequencing Unit |
| 225117 | *Pyrus x bretschneideri* | NCBI |
| VITVI | *Vitis vinifera* | NCBI |
| CARPA | *Carica papaya* | NCBI |
| CITLA | *Citrullus lanatus* | Cucurbit Genomics Database |
| CUCME | *Cucumis melo* | melonomics.net |
| POPTR | *Populus trichocarpa* | EnsemblPlants - Release 15 |
| ARATH | *Arabidopsis thaliana* | Ensembl Plants - Release 17 |
| MALDO | *Malus x domestica* | JGI - PhytozomeV11 |
| PRUDU | *Prunus dulcis* | Prunus dulcis genome project |
| PRUVE | *Prunus persica* | JGI - PhytozomeV11 |
| SOYBN | *Glycine max* | Ensembl Plants - Release 17 |
| ORYSJ | *Oryza sativa subsp. japonica* | Ensembl Plants - Release 22 |
| PRUAV | *Prunus avium* | NCBI |

**Table S3**. Almond cultivars selected and their main characteristics.

| Cultivar | Origin | Pedigree | Self-incompatibility[a] | Shell hardness | Bloom time |
|---|---|---|---|---|---|
| **Aï** | France | unknown | SI | Semi-hard | Very early |
| **Belle d'Aurons** | France | unknown | SI | Semi-hard | Early |
| **Cristomorto** | Italy | unknown | SI | Hard | Late |
| **Desmayo largueta** | Spain | unknown | SI | Hard | Early |
| **Falsa Barese** | Italy | unknown | SC | Hard | Late |
| **Genco** | Italy | unknown | SC | Hard | Late |
| **Marcona** | Spain | unknown | SI | Hard | Mid |
| **Nonpareil** | U.S.A. | unknown | SI | soft | Mid |
| **Ripon** | U.S.A. | unknown | SI | soft | Very late |
| **Texas** | U.S.A. | Seedling from Languedoc | SI | Semi-hard | Late |
| **Vivot** | Spain | unknown | SI | Hard | Early |

[a] SI: self-incompatible; SC: self-compatible

**Table S4**. Mapping of SNP markers from the TxE linkage map onto the almond assembly

| Chromosome | SNPs in map[a] | SNPs in assembly[b] | SNPs in pseudo-molecules[c] | Syntenic and collinear markers[d] | Non-syntenic and non-collinear markers[e] | Non-collinear markers | Different LG (marker-LG) |
|---|---|---|---|---|---|---|---|
| **Pd01** | 248 | 232 | 231 | 229 | 3 | SNP_IGA_67137; SNP_IGA_67265 | SNP_IGA_96232 (Pd06) |
| **Pd02** | 290 | 237 | 237 | 234 | 3 | SNP_IGA_141858; SNP_IGA_158810; SNP_IGA_158824 | |
| **Pd03** | 197 | 168 | 163 | 160 | 8 | SNP_IGA_300877; SNP_IGA_300953; SNP_IGA_356407 | SNP_IGA_887061 (pdulcis26_s0827); SNP_IGA_330713 (pdulcis26_s0920); SNP_IGA_356484, SNP_IGA_356701 (pdulcis26_s0416); snp_3_21905073 (pdulcis26_s0407) |
| **Pd04** | 391 | 350 | 350 | 349 | 1 | SNP_IGA_406345 | |
| **Pd05** | 134 | 116 | 116 | 116 | 0 | | |
| **Pd06** | 239 | 217 | 215 | 213 | 4 | SNP_IGA_619081; SNP_IGA_694408 | SNP_IGA_9623 (Pp01); snp_6_21067422 (pdulcis26_s0817) |
| **Pd07** | 154 | 133 | 132 | 128 | 5 | SNP_IGA_722889; SNP_IGA_722899; SNP_IGA_722921; SNP_IGA_722928 | SNP_IGA_727690 (Pd08) |
| **Pd08** | 180 | 156 | 153 | 149 | 7 | SNP_IGA_809997;SNP_IGA_809892;SNP_IGA_809771;SNP_IGA_816287 | SNP_IGA_812523 (pdulcis26_s1221); SNP_IGA_827612 (pdulcis26_s0616); SNP_IGA_853053 (pdulcis26_s0722) |
| **Total** | **1833** | **1609** | **1597** | **1578** | **31** | | |

[a]number of TxE SNPs belonging to the corresponding peach linkage group
[b]number of TxE SNPs mapped onto the almond assembly
[c]number of TxE SNPs mapped onto the corresponding almond linkage group
[d]number of TxE SNPs that have the same order onto the corresponding almond linkage group
[e]number of TxE SNPs that are mapped on the corresponding linkage group but have a different order than the TxE ("order change") or TxE SNPs that map onto different linkage group/scaffolds ("different linkage group/scaffold")

**Table S5**. General statistics of TE annotation in *P. dulcis* and *P. persica*.

| | ***P. dulcis*** | ***P. persica*** |
|---|---|---|
| **Total TE coverage (%)** | 38.21 | 37.60 |
| **Number of consensus sequences** | 3,994 | 3,922 |
| **Consensuses with full-length copies** | 2,307 (57.7 %) | 2,116 (53.9 %) |
| **Number of copies** | 73,679 | 65,472 |
| **Number of full-length copies** | 7,966 (10.8 %) | 9,143 (14.0 %) |

**Table S6**. Percentage of TE coverage at the order level in *P. dulcis* and *P. persica*.

| TE class | TE order | *P. dulcis* | *P. persica* |
|---|---|---|---|
| **Class I** | LTR | 21.28 | 19.8 |
| | LINE | 1.74 | 1.93 |
| | SINE | 0.27 | 0.4 |
| | DIRS | 0.28 | 0.33 |
| **Class II** | TIR | 13 | 14.15 |
| | Helitron | 1.27 | 0.81 |
| | Maverick | 0.35 | 0.18 |

**Table S7**. Detailed annotation of LTR retrotransposons and MITEs in *P. dulcis* and *P. persica.*

| **LTR-retrotransposon superfamily** | ***P. dulcis*** | ***P. persica*** |
|---|---|---|
| **Copia** | 964 | 1,040 |
| **Gypsy** | 392 | 517 |
| **Unclassified*** | 792 | 658 |
| **TOTAL LTR-retrotransposons** | 2,148 | 2,215 |
| MITES | | |
| **Full length** | 10,460 | 8,738 |
| **Partial MITE copies** | 56,196 | 53,711 |
| **TOTAL MITEs** | 66,656 | 62,449 |

*Elements carrying Long Terminal Repeats but lacking one or more coding domains

**Table S8.** Estimated dates (Mya) and 95% highest posterior density (HPD). Node numbers correspond to nodes in Figure 2A.

| **Node** | **meandate** | **stderr** | **inf95** | **sup95** |
|---|---|---|---|---|
| 1 | 5.8784 | 8.17344 | 0.737743 | 32.0312 |
| 2 | 20.8363 | 19.4593 | 2.31114 | 65.1583 |
| 3 | 62.0399 | 8.63017 | 49.329 | 83.1513 |
| 4 | 81.5 | 11.5929 | 60.975 | 108.329 |
| 5 | 23.6605 | 17.3587 | 4.12562 | 69.2511 |
| 6 | 97.5383 | 10.5984 | 81.064 | 123.354 |
| 7 | 126.924 | 14.4311 | 103.573 | 158.422 |
| 8 | 114.157 | 16.593 | 83.9286 | 149.426 |
| 9 | 25.331 | 16.5652 | 5.74043 | 68.7239 |
| 10 | 137.574 | 15.3032 | 112.234 | 171.606 |
| 11 | 116.867 | 20.3396 | 74.1546 | 156.087 |
| 12 | 81.7086 | 22.8562 | 39.3539 | 127.633 |
| 13 | 150.469 | 16.6399 | 122.542 | 186.548 |
| 14 | 156.15 | 17.7069 | 125.883 | 193.866 |
| 15 | 201.166 | 21.9517 | 162.337 | 248.811 |
| 16 | 220.6 | 22.4862 | 180.6 | 267.057 |

**Table S9.** List of the GO terms enriched in protein families of almond and peach that duplicated at the last common ancestor of *Prunus* species. The columns show, in this order: the GO term category, GO term, term level, enrichment *p* value, and term name.

| | | | | |
|---|---|---|---|---|
| ***P. dulcis*** | | | | |
| Term category | term | term level | adj.pvalue | term name |
| molecular_function | GO:0004812 | 1 | 9.25E-04 | aminoacyl-tRNA ligase activity |
| molecular_function | GO:0005506 | 1 | 5.76E-20 | iron ion binding |
| molecular_function | GO:0008168 | 1 | 3.65E-04 | methyltransferase activity |
| molecular_function | GO:0008171 | 1 | 6.52E-10 | O-methyltransferase activity |
| molecular_function | GO:0016705 | 1 | 3.15E-20 | oxidoreductase activity, acting on paired donors, with incorporation or reduction of molecular oxygen |
| molecular_function | GO:0016747 | 1 | 1.45E-12 | transferase activity, transferring acyl groups other than amino-acyl groups |
| molecular_function | GO:0016758 | 1 | 1.38E-05 | transferase activity, transferring hexosyl groups |
| molecular_function | GO:0020037 | 1 | 3.03E-16 | heme binding |
| biological_process | GO:0006418 | 1 | 7.65E-04 | tRNA aminoacylation for protein translation |
| biological_process | GO:0055114 | 1 | 8.35E-16 | oxidation-reduction process |
| ***P. persica*** | | | | |
| Term category | term | term level | adj.pvalue | term name |
| molecular_function | GO:0004812 | 1 | 3.65E-04 | aminoacyl-tRNA ligase activity |
| molecular_function | GO:0004970 | 1 | 1.53E-06 | ionotropic glutamate receptor activity |
| molecular_function | GO:0005216 | 1 | 1.11E-08 | ion channel activity |
| molecular_function | GO:0005506 | 1 | 7.50E-38 | iron ion binding |
| molecular_function | GO:0010333 | 1 | 3.81E-08 | terpene synthase activity |
| molecular_function | GO:0016491 | 1 | 6.24E-11 | oxidoreductase activity |
| molecular_function | GO:0016705 | 1 | 1.72E-39 | oxidoreductase activity, acting on paired donors, with incorporation or reduction of molecular oxygen |
| molecular_function | GO:0016758 | 1 | 3.28E-04 | transferase activity, transferring hexosyl groups |
| molecular_function | GO:0016829 | 1 | 1.15E-07 | lyase activity |
| molecular_function | GO:0020037 | 1 | 1.62E-35 | heme binding |
| molecular_function | GO:0030145 | 1 | 1.68E-05 | manganese ion binding |
| molecular_function | GO:0033926 | 1 | 9.02E-05 | glycopeptide alpha-N- |

| | | | | |
|---|---|---|---|---|
| | | | | acetylgalactosaminidase activity |
| molecular_function | GO:0045735 | 1 | 5.06E-05 | nutrient reservoir activity |
| molecular_function | GO:0071949 | 1 | 1.20E-05 | FAD binding |
| biological_process | GO:0006418 | 1 | 4.30E-04 | tRNA aminoacylation for protein translation |
| biological_process | GO:0006811 | 1 | 1.78E-08 | ion transport |
| biological_process | GO:0008152 | 1 | 3.45E-06 | metabolic process |
| biological_process | GO:0009733 | 1 | 2.10E-05 | response to auxin |
| biological_process | GO:0055114 | 1 | 3.18E-31 | oxidation-reduction process |

**Table S10.** List of the GO terms enriched in the protein families lost specifically in peach and almond. The columns show in this order: the term category, the GO term, the term level, the p value, and the term name.

| ***P. persica*** **losses** | | | | |
|---|---|---|---|---|
| Term category | term | term level | adj.pvalue | term name |
| molecular_function | GO:0003899 | 1 | 4.95E-05 | DNA-directed 5'-3' RNA polymerase activity |
| molecular_function | GO:0016747 | 1 | 3.01E-07 | transferase activity, transferring acyl groups other than amino-acyl groups |
| cellular_component | GO:0009507 | 1 | 4.95E-05 | chloroplast |
| biological_process | GO:0006351 | 1 | 4.72E-04 | transcription, DNA-templated |
| biological_process | GO:0015986 | 1 | 4.72E-04 | ATP synthesis coupled proton transport |
| ***P. dulcis*** **losses** | | | | |
| Term category | term | term level | adj.pvalue | term name |
| molecular_function | GO:0003824 | 1 | 9.26E-04 | catalytic activity |
| molecular_function | GO:0004867 | 1 | 3.56E-04 | serine-type endopeptidase inhibitor activity |
| molecular_function | GO:0004970 | 1 | 8.81E-09 | ionotropic glutamate receptor activity |
| molecular_function | GO:0005216 | 1 | 1.10E-04 | ion channel activity |
| molecular_function | GO:0005506 | 1 | 1.03E-25 | iron ion binding |
| molecular_function | GO:0016491 | 1 | 6.07E-04 | oxidoreductase activity |
| molecular_function | GO:0016705 | 1 | 2.50E-27 | oxidoreductase activity, acting on paired donors, with incorporation or reduction of molecular oxygen |
| molecular_function | GO:0016887 | 1 | 2.63E-04 | ATPase activity |
| molecular_function | GO:0020037 | 1 | 3.89E-27 | heme binding |
| molecular_function | GO:0045735 | 1 | 5.92E-04 | nutrient reservoir activity |
| molecular_function | GO:0071949 | 1 | 1.35E-04 | FAD binding |
| cellular_component | GO:0016020 | 1 | 1.80E-06 | membrane |
| biological_process | GO:0006811 | 1 | 1.35E-04 | ion transport |
| biological_process | GO:0006869 | 1 | 5.14E-04 | lipid transport |
| biological_process | GO:0009733 | 1 | 2.23E-13 | response to auxin |
| biological_process | GO:0055114 | 1 | 3.20E-22 | oxidation-reduction process |

**Table S11**. Mapping statistics for the resequenced almond cultivars

| Cultivar | Total reads | Reads mapped | % reads mapped | Average depth | Median depth | % genome coverage (≥5 reads) |
|---|---|---|---|---|---|---|
| **Aï** | 90.435.308 | 84.889.166 | 93 | 43,73 | 32,00 | 93 |
| **Belle d'Aurons** | 88.461.982 | 83.002.863 | 93 | 42,74 | 31,00 | 94 |
| **Cristomorto** | 73.876.698 | 68.824.743 | 93 | 36,04 | 26,00 | 92 |
| **Desmayo Largueta** | 91.447.444 | 85.201.669 | 93 | 43,90 | 32,00 | 94 |
| **Falsa Barese** | 81.923.166 | 64.775.811 | 79 | 27,68 | 19,00 | 90 |
| **Genco** | 86.412.808 | 69.105.067 | 79 | 29,67 | 21,00 | 91 |
| **Marcona** | 107.453.860 | 100.455.135 | 93 | 51,43 | 39,00 | 94 |
| **Nonpareil** | 118.234.470 | 111.519.612 | 94 | 47,02 | 36,00 | 95 |
| **Ripon** | 93.785.436 | 76.095.528 | 81 | 32,37 | 23,00 | 94 |
| **Vivot** | 86.988.642 | 82.044.847 | 94 | 42,83 | 31,00 | 93 |

**Table S12.** Variant distribution across the almond pseudomolecules

| Chromo-some | Aï | | Belle d'Aurons | | Crstomorto | | Desmayo largueta | | Falsa Barese | | Genco | | Marcona | | Nonpareil | | Ripon | | Vivot | | Total | |
|---|---|---|---|---|---|---|---|---|---|---|---|---|---|---|---|---|---|---|---|---|---|---|
| | SNPs | INDELs | SNPs | INDELs | SNPs | INDELs | SNPs | INDELs | SNPs | INDELs | SNPs | INDELs | SNPs | INDELs | SNPs | INDELs | SNPs | INDELs | SNPs | INDELs | SNPs | INDELs |
| **Pd01** | 195.744 | 27.391 | 192.338 | 26.158 | 178.882 | 24.267 | 189.620 | 26.269 | 153.651 | 16.244 | 177.310 | 19.320 | 200.737 | 28.342 | 217.220 | 29.385 | 159.769 | 18.616 | 181.801 | 25.188 | 419.236 | 65.221 |
| **Pd02** | 124.980 | 16.908 | 133.963 | 17.652 | 111.590 | 14.725 | 128.962 | 17.090 | 117.283 | 12.161 | 123.631 | 13.122 | 133.044 | 18.196 | 137.868 | 18.168 | 106.156 | 12.052 | 124.553 | 16.719 | 291.354 | 43.476 |
| **Pd03** | 122.012 | 16.100 | 121.631 | 15.825 | 112.409 | 14.564 | 113.649 | 14.858 | 107.302 | 11.207 | 110.401 | 11.704 | 119.119 | 16.025 | 132.760 | 17.096 | 102.613 | 11.292 | 114.244 | 14.872 | 257.739 | 38.013 |
| **Pd04** | 117.533 | 15.163 | 119.899 | 15.653 | 116.139 | 14.605 | 113.879 | 14.795 | 108.292 | 10.959 | 112.335 | 11.393 | 114.720 | 15.258 | 111.697 | 14.420 | 90.142 | 9.767 | 104.974 | 13.871 | 255.126 | 36.865 |
| **Pd05** | 86.423 | 12.270 | 84.640 | 11.718 | 84.973 | 11.324 | 87.550 | 12.256 | 85.256 | 9.348 | 79.821 | 8.869 | 87.923 | 12.464 | 99.811 | 13.938 | 75.648 | 9.144 | 88.430 | 12.185 | 194.446 | 30.704 |
| **Pd06** | 138.360 | 18.782 | 138.082 | 18.613 | 122.865 | 16.121 | 135.722 | 18.075 | 122.666 | 12.794 | 123.970 | 13.065 | 129.158 | 18.102 | 151.091 | 19.931 | 117.096 | 13.233 | 134.370 | 17.733 | 298.716 | 45.013 |
| **Pd07** | 102.611 | 14.033 | 112.403 | 14.773 | 97.897 | 12.946 | 100.450 | 13.503 | 85.230 | 8.571 | 100.484 | 10.408 | 99.824 | 14.153 | 97.094 | 13.346 | 86.007 | 9.760 | 106.521 | 14.292 | 231.097 | 34.542 |
| **Pd08** | 106.511 | 14.454 | 106.297 | 14.342 | 101.401 | 13.269 | 96.351 | 12.924 | 100.206 | 10.379 | 100.522 | 10.679 | 104.484 | 14.461 | 113.319 | 14.890 | 79.686 | 9.487 | 97.146 | 13.090 | 227.841 | 34.417 |
| **Un-anchored** | 10.683 | 927 | 12.959 | 1.107 | 8.911 | 778 | 10.610 | 885 | 9.946 | 612 | 11.272 | 709 | 10.761 | 947 | 11.899 | 968 | 10.280 | 719 | 10.472 | 885 | 28.027 | 2.544 |
| **Total** | 1.004.857 | 136.028 | 1.022.212 | 135.841 | 935.067 | 122.599 | 976.793 | 130.655 | 889.832 | 92.275 | 939.746 | 99.269 | 999.770 | 137.948 | 1.072.759 | 142.142 | 827.397 | 94.070 | 962.511 | 128.835 | 2.203.582 | 330.795 |

**Table S13**: Comparison of SNP variability parameters in *Prunus* species with whole genome sequences available.

| **Parameters** | **cherry** | **peach** | **almond** | **peach** | **almond** | **almond** |
|---|---|---|---|---|---|---|
| **Reference** | **Shirasawa et al. (2017)** | **Yu et al. (2018)** | | **Velasco et al. (2016)** | | **This paper** |
| **genome assembly (Mbp)** | 272.3 | 227,411,381 | 227,411,381[a] | - | - | 227,599,157 |
| **# samples** | 6 | 22 | 15 | 13 | 13 | 10 |
| **Average density (SNP/kbp)** | 2.5 | 2.1 | 19.1 | - | - | 6.2[b] |
| **Average # of heterozygous** | 463,240.5 | 274,037.9 | 1.331.604,7 | - | - | 995,912.4[c] |
| **Average heterozygosity (%)** | 0.170 | 0.121 | 0.685 | - | - | 0.438 |
| **Average Inbreeding coefficient** | - | 0.3192 | -0.0909 | 0.197 | 0.002 | -0.0017 |

[a] Yu et al. used the peach genome as reference sequence, for both peach and almond.

[b] Lower SNP density, as compared to Yu et al. (2018) is due to the use of the almond genome as a reference in this paper vs. the pach genoms in Yu's, resulting in the inclusion of homozygous almond SNPs versus the peach sequence. Usage of the pdulcis26 assembly with our data de facto excludes these homozygous SNPs from our analysis.

[c] This number corresponds to ~⅔ of the heterozygous SNPs found by Yu et al. (1,331,604), probably due to the smaller number of almond varieties used in our analysis and/or lower inherent heterozygosity of the selected varieties.

**Table S14.** Deletions in ten almond and one peach cultivars compared to the almond reference sequence and deletions that contain transposable element (TE) sequences.

| | 1-20 | | | 21-50 | | | 51-500 | | | 501-10000 | | | 10001-50000 | | | Large deletions (>50bp) | | | Total | | |
|---|---|---|---|---|---|---|---|---|---|---|---|---|---|---|---|---|---|---|---|---|---|
| **Line** | **all** | **TEs** | **%** | **all** | **TEs** | **%** | **all** | **TEs** | **%** | **all** | **TEs** | **%** | **all** | **TEs** | **%** | **all** | **TEs** | **%** | **all** | **TEs** | **%** |
| **ALMOND** | | | | | | | | | | | | | | | | | | | | | |
| Aï | 65,431 | 17,578 | 26,9 | 3,445 | 914 | 26.5 | 114 | 31 | 27.2 | 30 | 29 | 96.7 | 3 | 3 | 100.0 | 147 | 63 | 42.9 | 69,023 | 18,555 | 26.9 |
| Belle d'Aurons | 65,739 | 18,096 | 27,5 | 3,387 | 924 | 27.3 | 96 | 27 | 28.1 | 22 | 21 | 95.5 | 3 | 3 | 100.0 | 121 | 51 | 42.1 | 69,247 | 19,071 | 27.5 |
| Cristomorto | 59,715 | 15,423 | 25,8 | 2,792 | 702 | 25.1 | 52 | 12 | 23.1 | 13 | 12 | 92.3 | 2 | 2 | 100.0 | 67 | 26 | 38.8 | 62,574 | 16,151 | 25.8 |
| Desmayo Largueta | 63,032 | 16,735 | 26,6 | 3,135 | 835 | 26.6 | 116 | 33 | 28.4 | 43 | 38 | 88.4 | 5 | 5 | 100.0 | 164 | 76 | 46.3 | 66,331 | 17,646 | 26.6 |
| Falsa Barese | 46,450 | 11,767 | 25,3 | 1,483 | 412 | 27.8 | 18 | 6 | 33.3 | 4 | 4 | 100.0 | - | - | - | 22 | 10 | 45.5 | 47,955 | 12,189 | 25.4 |
| Genco | 49,842 | 12,530 | 25,1 | 1,739 | 436 | 25.1 | 18 | 4 | 22.2 | 7 | 7 | 100.0 | - | - | - | 25 | 11 | 44.0 | 51,606 | 12,977 | 25.1 |
| Marcona | 66,438 | 18,136 | 27,3 | 3,378 | 924 | 27.4 | 219 | 68 | 31.1 | 74 | 72 | 97.3 | 8 | 8 | 100.0 | 301 | 148 | 49.2 | 70,117 | 19,208 | 27.4 |
| Nonpareil | 69,237 | 19,304 | 27,9 | 2,785 | 791 | 28.4 | 151 | 65 | 43.0 | 63 | 56 | 88.9 | 5 | 5 | 100.0 | 219 | 126 | 57.5 | 72,241 | 20,221 | 28.0 |
| Ripon | 46,164 | 12,016 | 26,0 | 1,799 | 487 | 27.1 | 12 | 5 | 41.7 | 5 | 5 | 100.0 | - | - | - | 17 | 10 | 58.8 | 47,980 | 12,513 | 26.1 |
| Vivot | 62,104 | 16,556 | 26,7 | 3,065 | 805 | 26.3 | 96 | 32 | 33.3 | 38 | 33 | 86.8 | 2 | 2 | 100.0 | 136 | 67 | 49.3 | 65,305 | 17,428 | 26.7 |
| **Total** | 594,152 | 158,141 | 26.6 | 27,008 | 7,230 | 26.8 | 892 | 283 | 31.7 | 299 | 277 | 92.6 | 28 | 28 | 100.0 | 1,219 | 588 | 48.2 | 622,379 | 17,429 | 26.7 |
| **Average** | 59,415 | 15,814 | 26,6 | 2,701 | 723 | 26.8 | 88 | 28 | 31.7 | 30 | 28 | 92.6 | 3 | 3 | 100.0 | 120 | 59 | 49.0 | 62,236 | 17,430 | 26.7 |
| **PEACH** | | | | | | | | | | | | | | | | | | | | | |
| Earlygold | 120,418 | 19,778 | 16,4 | 4,283 | 609 | 14.2 | 1,090 | 190 | 17.4 | 332 | 274 | 82.5 | 14 | 14 | 100,0 | 1,436 | 478 | 33.3 | 126,137 | 12,513 | 9.9 |

**Table S15**. Summary of variants detected between *P. dulcis* and *P. persica* assemblies.

| | Insertion | | | Deletion | | | Repeat_expansion | | | Repeat_contraction | | |
|---|---|---|---|---|---|---|---|---|---|---|---|---|
| **Size range** | **Count** | **Total bp** | **TE related** | **Count** | **Total bp** | **TE related** | **Count** | **Total bp** | **TE related** | **Count** | **Total bp** | **TE related** |
| 20-50 bp * | 5,945 | 170,266 | N/A | 5723 | 163,874 | N/A | 79 | 2,576 | N/A | 65 | 2,276 | N/A |
| 50-500 bp | 1,644 | 223,420 | 592 | 1630 | 244,546 | 317 | 571 | 133,996 | 122 | 666 | 160,939 | 137 |
| 500-10000 bp | 472 | 947,624 | 358 | 497 | 1,204,612 | 299 | 1,099 | 2,990,754 | 266 | 1550 | 5,096,489 | 293 |
| 10000-50000 bp | 9 | 104,033 | 4 | 41 | 574,692 | 25 | 52 | 676,160 | 10 | 337 | 5,227,362 | 60 |
| Total | 8,070 | 1,445,343 | 954 | 7,929 | 2,189,624 | 641 | 1,801 | 3,803,486 | 398 | 2,618 | 10,487,066 | 490 |

* This size interval does not match with the size of any transposon family

**Table S16**. List of the 97 genes potentially involved in mesocarp development. (See Excel file Table S16)

**TAIR_name_description (Best hit)**

Symbols: WUS, PGA6, WUS1 | Homeodomain-like superfamily protein | chr2:7809100-7810671 REVERSE LENGTH=292
Homeodomain-like superfamily protein
Symbols: WOX1 | WUSCHEL related homeobox 1 | chr3:6161155-6163183 REVERSE LENGTH=350
WUSCHEL related homeobox 1
Symbols: WOX2 | WUSCHEL related homeobox 2 | chr5:23933408-23934627 REVERSE LENGTH=260
WUSCHEL related homeobox 2
Symbols: PRS, WOX3, PRS1 | Homeodomain-like superfamily protein | chr2:12262115-12263286 FORWARD LENGTH=244
Homeodomain-like superfamily protein
Symbols: WOX4 | WUSCHEL related homeobox 4 | chr1:17236903-17237953 REVERSE LENGTH=251
WUSCHEL related homeobox 4
Symbols: WOX5 | WUSCHEL related homeobox 5 | chr3:3527606-3528263 FORWARD LENGTH=182
WUSCHEL related homeobox 5
Symbols: WOX9, HB-3, STIP | homeobox-3 | chr2:14341639-14343597 REVERSE LENGTH=378
homeobox-3
Symbols: WOX11 | WUSCHEL related homeobox 11 | chr3:889515-892162 REVERSE LENGTH=268
WUSCHEL related homeobox 11
Symbols: HB-4, WOX13, ATWOX13 | WUSCHEL related homeobox 13 | chr4:16875814-16877167 REVERSE LENGTH=268
WUSCHEL related homeobox 13
Symbols: HB-4, WOX13, ATWOX13 | WUSCHEL related homeobox 13 | chr4:16875814-16877167 REVERSE LENGTH=268
WUSCHEL related homeobox 13
Symbols: STM, BUM1, SHL, WAM1, BUM, WAM | KNOX/ELK homeobox transcription factor | chr1:23058796-23061722 REVERSE LENGTH=382
KNOX/ELK homeobox transcription factor
Symbols: STM, BUM1, SHL, WAM1, BUM, WAM | KNOX/ELK homeobox transcription factor | chr1:23058796-23061722 REVERSE LENGTH=382
KNOX/ELK homeobox transcription factor
Symbols: KNAT1, BP, BP1 | KNOTTED-like from Arabidopsis thaliana | chr4:5147969-5150610 REVERSE LENGTH=398
KNOTTED-like from Arabidopsis thaliana
Symbols: KNAT3 | KNOTTED1-like homeobox gene 3 | chr5:8736208-8738115 FORWARD LENGTH=431
KNOTTED1-like homeobox gene 3

Symbols: KNAT3 | KNOTTED1-like homeobox gene 3 | chr5:8736208-8738115 FORWARD LENGTH=431
KNOTTED1-like homeobox gene 3
Symbols: KNAT6, KNAT6L, KNAT6S | KNOTTED1-like homeobox gene 6 | chr1:8297499-8302492 REVERSE LENGTH=327
KNOTTED1-like homeobox gene 6
Symbols: KNAT6, KNAT6L, KNAT6S | KNOTTED1-like homeobox gene 6 | chr1:8297499-8302492 REVERSE LENGTH=327
KNOTTED1-like homeobox gene 6
Symbols: KNAT6, KNAT6L, KNAT6S | KNOTTED1-like homeobox gene 6 | chr1:8297499-8302492 REVERSE LENGTH=327
KNOTTED1-like homeobox gene 6
Symbols: BEL1 | POX (plant homeobox) family protein | chr5:16580424-16583770 FORWARD LENGTH=611
POX (plant homeobox) family protein
Symbols: PHB, ATHB14, ATHB-14, PHB-1D | Homeobox-leucine zipper family protein / lipid-binding START domain-containing protein | chr2:14639548-14643993 REVERSE LENG
Homeobox-leucine zipper family protein / lipid-binding START domain-containing protein
Symbols: REV, IFL, IFL1 | Homeobox-leucine zipper family protein / lipid-binding START domain-containing protein | chr5:24397734-24401933 FORWARD LENGTH=842
Homeobox-leucine zipper family protein / lipid-binding START domain-containing protein
Symbols: ATHB-15, ATHB15, CNA, ICU4 | Homeobox-leucine zipper family protein / lipid-binding START domain-containing protein | chr1:19409913-19413961 REVERSE LENGTH
Homeobox-leucine zipper family protein / lipid-binding START domain-containing protein
Symbols: AG | K-box region and MADS-box transcription factor family protein | chr4:10383917-10388272 FORWARD LENGTH=252
K-box region and MADS-box transcription factor family protein
K-box region and MADS-box transcription factor family protein
Symbols: SHP1, AGL1 | K-box region and MADS-box transcription factor family protein | chr3:21739150-21741766 FORWARD LENGTH=248
K-box region and MADS-box transcription factor family protein
K-box region and MADS-box transcription factor family protein
Symbols: SEP1, AGL2 | K-box region and MADS-box transcription factor family protein | chr5:5151594-5153767 REVERSE LENGTH=251
K-box region and MADS-box transcription factor family protein
Symbols: SEP1, AGL2 | K-box region and MADS-box transcription factor family protein | chr5:5151594-5153767 REVERSE LENGTH=251
K-box region and MADS-box transcription factor family protein
Symbols: SEP2, AGL4 | K-box region and MADS-box transcription factor family protein | chr3:464554-466687 REVERSE LENGTH=250
K-box region and MADS-box transcription factor family protein
Symbols: AGL8, FUL | AGAMOUS-like 8 | chr5:24502736-24506013 REVERSE LENGTH=242
AGAMOUS-like 8
Symbols: SEP3, AGL9 | K-box region and MADS-box transcription factor family protein | chr1:8593790-8595862 REVERSE LENGTH=250
K-box region and MADS-box transcription factor family protein
Symbols: STK, AGL11 | K-box region and MADS-box transcription factor family protein | chr4:6236713-6239409 REVERSE LENGTH=230
K-box region and MADS-box transcription factor family protein
Symbols: AP1, AGL7 | K-box region and MADS-box transcription factor family protein | chr1:25982576-25986102 REVERSE LENGTH=256
K-box region and MADS-box transcription factor family protein
Symbols: AP1, AGL7 | K-box region and MADS-box transcription factor family protein | chr1:25982576-25986102 REVERSE LENGTH=256
K-box region and MADS-box transcription factor family protein
Symbols: AP2 | Integrase-type DNA-binding superfamily protein | chr4:17400998-17403140 FORWARD LENGTH=432
Integrase-type DNA-binding superfamily protein

Symbols: AP3, ATAP3 | K-box region and MADS-box transcription factor family protein | chr3:20119428-20121087 REVERSE LENGTH=232
K-box region and MADS-box transcription factor family protein
Symbols: AP3, ATAP3 | K-box region and MADS-box transcription factor family protein | chr3:20119428-20121087 REVERSE LENGTH=232
K-box region and MADS-box transcription factor family protein
Symbols: PI | K-box region and MADS-box transcription factor family protein | chr5:6829203-6831208 FORWARD LENGTH=208
K-box region and MADS-box transcription factor family protein
Symbols: ER, QRP1 | Leucine-rich receptor-like protein kinase family protein | chr2:11208367-11213895 REVERSE LENGTH=976
Leucine-rich receptor-like protein kinase family protein
Symbols: BAM1 | Leucine-rich receptor-like protein kinase family protein | chr5:26281826-26284945 FORWARD LENGTH=1003
Leucine-rich receptor-like protein kinase family protein
Symbols: BAM1 | Leucine-rich receptor-like protein kinase family protein | chr5:26281826-26284945 FORWARD LENGTH=1003
Leucine-rich receptor-like protein kinase family protein
Symbols: BAM1 | Leucine-rich receptor-like protein kinase family protein | chr5:26281826-26284945 FORWARD LENGTH=1003
Leucine-rich receptor-like protein kinase family protein
Symbols: BAM3 | Leucine-rich receptor-like protein kinase family protein | chr4:10949822-10952924 FORWARD LENGTH=992
Leucine-rich receptor-like protein kinase family protein
Symbols: CLE12 | CLAVATA3/ESR-RELATED 12 | chr1:25841079-25841435 REVERSE LENGTH=118
CLAVATA3/ESR-RELATED 12
Symbols: CLE13 | CLAVATA3/ESR-RELATED 13 | chr1:27815822-27816145 FORWARD LENGTH=107
CLAVATA3/ESR-RELATED 13
Symbols: CLE25 | CLAVATA3/ESR-RELATED 25 | chr3:10670220-10670931 REVERSE LENGTH=81
CLAVATA3/ESR-RELATED 25
Symbols: CLV2, AtRLP10 | Leucine-rich repeat (LRR) family protein | chr1:24286943-24289105 FORWARD LENGTH=720
Leucine-rich repeat (LRR) family protein
Symbols: CLV1, FAS3, FLO5, ATCLV1 | Leucine-rich receptor-like protein kinase family protein | chr1:28463631-28466652 REVERSE LENGTH=980
Leucine-rich receptor-like protein kinase family protein
Symbols: CLV1, FAS3, FLO5, ATCLV1 | Leucine-rich receptor-like protein kinase family protein | chr1:28463631-28466652 REVERSE LENGTH=980
Leucine-rich receptor-like protein kinase family protein
Symbols: RPK2, TOAD2, CLI1 | receptor-like protein kinase 2 | chr3:380726-384181 FORWARD LENGTH=1151
receptor-like protein kinase 2
Symbols: RPK2, TOAD2, CLI1 | receptor-like protein kinase 2 | chr3:380726-384181 FORWARD LENGTH=1151
receptor-like protein kinase 2
Symbols: HEC1 | basic helix-loop-helix (bHLH) DNA-binding superfamily protein | chr5:26766276-26767001 FORWARD LENGTH=241
basic helix-loop-helix (bHLH) DNA-binding superfamily protein
Symbols: HEC2 | basic helix-loop-helix (bHLH) DNA-binding superfamily protein | chr3:18657423-18658118 REVERSE LENGTH=231
basic helix-loop-helix (bHLH) DNA-binding superfamily protein
Symbols: IND1, GT140, IND, EDA33 | basic helix-loop-helix (bHLH) DNA-binding superfamily protein | chr4:42601-43197 REVERSE LENGTH=198
basic helix-loop-helix (bHLH) DNA-binding superfamily protein
Symbols: SPT | basic helix-loop-helix (bHLH) DNA-binding superfamily protein | chr4:17414167-17415945 FORWARD LENGTH=373
basic helix-loop-helix (bHLH) DNA-binding superfamily protein

Symbols: SPT | basic helix-loop-helix (bHLH) DNA-binding superfamily protein | chr4:17414167-17415945 FORWARD LENGTH=373
basic helix-loop-helix (bHLH) DNA-binding superfamily protein
Symbols: ARF1 | auxin response factor 1 | chr1:21980414-21984193 FORWARD LENGTH=660
auxin response factor 1
Symbols: ARF1 | auxin response factor 1 | chr1:21980414-21984193 FORWARD LENGTH=665
auxin response factor 9
Symbols: ARF1 | auxin response factor 1 | chr1:21980414-21984193 FORWARD LENGTH=662
auxin response factor 1
Symbols: ARF2, ARF1-BP, HSS, ORE14 | auxin response factor 2 | chr5:24910859-24914680 FORWARD LENGTH=859
auxin response factor 2
Symbols: ETT, ARF3 | Transcriptional factor B3 family protein / auxin-responsive factor AUX/IAA-related | chr2:14325444-14328613 REVERSE LENGTH=608
Transcriptional factor B3 family protein / auxin-responsive factor AUX/IAA-related
Symbols: ARF4 | auxin response factor 4 | chr5:24308558-24312187 REVERSE LENGTH=788
auxin response factor 4
Symbols: MP, ARF5, IAA24 | Transcriptional factor B3 family protein / auxin-responsive factor AUX/IAA-related | chr1:6887353-6891182 FORWARD LENGTH=902
Transcriptional factor B3 family protein / auxin-responsive factor AUX/IAA-related
Symbols: ARF6 | auxin response factor 6 | chr1:10686125-10690036 REVERSE LENGTH=933
auxin response factor 6
Symbols: ARF6 | auxin response factor 6 | chr1:10686125-10690036 REVERSE LENGTH=933
auxin response factor 6
Symbols: ARF8, ATARF8 | auxin response factor 8 | chr5:14630151-14634106 FORWARD LENGTH=811
auxin response factor 8
Symbols: ARF9 | auxin response factor 9 | chr4:12451592-12454737 FORWARD LENGTH=638
auxin response factor 9
Symbols: ARF10 | auxin response factor 10 | chr2:12114331-12116665 FORWARD LENGTH=693
auxin response factor 10
Symbols: ARF16 | auxin response factor 16 | chr4:14703369-14705564 REVERSE LENGTH=670
auxin response factor 16
Symbols: ARF16 | auxin response factor 16 | chr4:14703369-14705564 REVERSE LENGTH=670
auxin response factor 16
Symbols: ARF17 | auxin response factor 17 | chr1:29272405-29275193 FORWARD LENGTH=585
auxin response factor 17
Symbols: ARF19, IAA22, ARF11 | auxin response factor 19 | chr1:6628395-6632779 REVERSE LENGTH=1086
auxin response factor 19
Symbols: ARF19, IAA22, ARF11 | auxin response factor 19 | chr1:6628395-6632779 REVERSE LENGTH=1086
auxin response factor 19
Symbols: ATOFP4, OFP4 | ovate family protein 4 | chr1:2124854-2125801 REVERSE LENGTH=315
ovate family protein 4
Symbols: ATOFP5, OFP5 | ovate family protein 5 | chr4:10337449-10338498 FORWARD LENGTH=349
ovate family protein 5

Symbols: ATOFP5, OFP5 | ovate family protein 5 | chr4:10337449-10338498 FORWARD LENGTH=349
ovate family protein 5
Symbols: AGO1 | Stabilizer of iron transporter SufD / Polynucleotidyl transferase | chr1:17886285-17891892 REVERSE LENGTH=1048
Stabilizer of iron transporter SufD / Polynucleotidyl transferase
Symbols: AGO1 | Stabilizer of iron transporter SufD / Polynucleotidyl transferase | chr1:17886285-17891892 REVERSE LENGTH=1048
Stabilizer of iron transporter SufD / Polynucleotidyl transferase
Symbols: ZLL, AGO10 | Stabilizer of iron transporter SufD / Polynucleotidyl transferase | chr5:17611939-17616562 FORWARD LENGTH=988
Stabilizer of iron transporter SufD / Polynucleotidyl transferase
Symbols: CYCA2;1 | cyclin a2;1 | chr5:8815230-8817566 FORWARD LENGTH=437
cyclin a2;1
Symbols: CYCA2;4 | Cyclin A2;4 | chr1:30214694-30216861 FORWARD LENGTH=461
Cyclin A2;4
Cyclin A2;4
Symbols: HAM3, ATHAM3, LOM3 | GRAS family transcription factor | chr4:57429-59105 REVERSE LENGTH=558
GRAS family transcription factor
Symbols: HAM3, ATHAM3, LOM3 | GRAS family transcription factor | chr4:57429-59105 REVERSE LENGTH=558
GRAS family transcription factor
Symbols: HAM4 | GRAS family transcription factor | chr4:17306060-17307520 FORWARD LENGTH=486
GRAS family transcription factor
Symbols: NIK1 | NSP-interacting kinase 1 | chr5:5224264-5227003 FORWARD LENGTH=638
NSP-interacting kinase 1
Symbols: NIK1 | NSP-interacting kinase 1 | chr5:5224264-5227003 FORWARD LENGTH=638
NSP-interacting kinase 1
Symbols: NIK3 | NSP-interacting kinase 3 | chr1:22383601-22386931 REVERSE LENGTH=632
NSP-interacting kinase 3
Symbols: PIN1, ATPIN1 | Auxin efflux carrier family protein | chr1:27659772-27662876 FORWARD LENGTH=622
Auxin efflux carrier family protein
Symbols: PIN1, ATPIN1 | Auxin efflux carrier family protein | chr1:27659772-27662876 FORWARD LENGTH=622
Auxin efflux carrier family protein
Symbols: ULCS1, LC | Transducin/WD40 repeat-like superfamily protein | chr5:26466348-26468201 FORWARD LENGTH=331
Transducin/WD40 repeat-like superfamily protein
Symbols: CRC | Plant-specific transcription factor YABBY family protein | chr1:26007734-26008940 REVERSE LENGTH=181
Plant-specific transcription factor YABBY family protein
Symbols: PID, ABR | Protein kinase superfamily protein | chr2:14589934-14591557 REVERSE LENGTH=438
Protein kinase superfamily protein
Symbols: FTA, PLP, ATFTA, PFT/PGGT-IALPHA | farnesyltransferase A | chr3:21944209-21945781 FORWARD LENGTH=326
farnesyltransferase A
Symbols: ULT1, ULT | Developmental regulator, ULTRAPETALA | chr4:13985753-13987050 FORWARD LENGTH=237
Developmental regulator, ULTRAPETALA
Symbols: SOL2, CRN | Protein kinase superfamily protein | chr5:4252924-4254215 REVERSE LENGTH=401

Protein kinase superfamily protein
Symbols: KAPP, RAG1 | kinase associated protein phosphatase | chr5:6488450-6493182 FORWARD LENGTH=581
kinase associated protein phosphatase
Symbols: TSL | Protein kinase superfamily protein | chr5:7098213-7102970 FORWARD LENGTH=688
Protein kinase superfamily protein
Symbols: AXR1 | NAD(P)-binding Rossmann-fold superfamily protein | chr1:1498357-1501775 REVERSE LENGTH=540
NAD(P)-binding Rossmann-fold superfamily protein
Symbols: ATBARD1, BARD1 | breast cancer associated RING 1 | chr1:1036610-1040045 FORWARD LENGTH=713
breast cancer associated RING 1

**Table S17.** Methylation status on genes potentially involved in mesocarp development and presenting TE insertions in peach or almond

| | | | | | Gene | | | | | TE | | | | | |
|---|---|---|---|---|---|---|---|---|---|---|---|---|---|---|---|
| | | TAIR best hit | Species | Chromosome | Accession | size (bp) | CG context (%) | CHG context (%) | CHH context (%) | type of TE | size (bp) | CG context (%) | CHG context (%) | CHH context (%) | Position inside the gene /Distance to gene (bp) |
| **TE insertion in *P.dulcis*** | **Inside the gene** | **Auxin Response Factor 8** (ARF8) | Almond | Pd03 | Prudul26A012714 | 12264 | 71,43 | 1,23 | 0,21 | MITE | 197 | 95,69 | 48,11 | 4,56 | Intron 8 |
| | | | Peach | Pp03 | Prupe.3G011800 | 11090 | 80,81 | 1,12 | 0,06 | | | | | | |
| | **Upstream region of the gene** | **WUSCHEL related homeobox 11** (WOX11) | Almond | Pd07 | Prudul26A026968 | 2225 | 11,48 | 3,2 | 0,4 | MITE | 110 | 78,67 | 2,17 | 0 | 485 |
| | | | Peach | Pp07 | Prupe.7G016600 | 2622 | 14,09 | 2,91 | 0,58 | | | | | | |
| | | **Repressor of WUSCHEL1** (ROW1) | Almond | Pd08 | Prudul26A000820 | 4610 | 17,61 | 0,07 | 0,06 | MITE | 186 | 82,59 | 58,55 | 50,65 | 305 |
| | | | Peach | Pp08 | Prupe.8G218500 | 4891 | 18,91 | 0,08 | 0,02 | | | | | | |
| **TE insertion in *P. persica*** | **Upstream region of the gene** | **FruitFull/AGAMOUS-like 8** (FUL/AGL8) | Peach | Pp05 | Prupe.5G208500 | 3324 | 3,43 | 0,16 | 0,13 | MITE | 595 | 93,84 | 52,5 | 16,06 | 199 |
| | | | Almond | Pd05 | Prudul26A029983 | 2487 | 0 | 0 | 0,02 | | | | | | |
| | | **Tousled** (TSL) | Peach | Pp06 | Prupe.6G149600 | 9594 | 66,83 | 0,07 | 0,06 | DNA TE | 3164 | 93,91 | 65,74 | 10,83 | 267 |
| | | | Almond | Pd06 | Prudul26A019759 | 9641 | 59,58 | 0,38 | 0,04 | | | | | | |
| | | **WUSCHEL related homeobox 11** (WOX11) | Peach | Pp07 | Prupe.7G016600 | 2622 | 14,09 | 2,91 | 0,58 | MITE | 175 | 96,86 | 65,38 | 29,9 | 157 |
| | | | Almond | Pd07 | Prudul26A026968 | 2225 | 11,48 | 3,2 | 0,4 | | | | | | |
| | | **Auxin Response Factor 1** (ARF1) | Peach | Pp08 | Prupe.8G252300 | 6799 | 14,08 | 0,18 | 0,03 | MITE | 596 | 96,31 | 34,49 | 19,54 | 1061 |
| | | | Almond | Pd08 | Prudul26A013138 | 6645 | 2,93 | 0,06 | 0,02 | | | | | | |
